# Novel transcriptional activator TAC3 regulates age-dependent floral transition in Chinese fir (*Cunninghamia lanceolata*)

**DOI:** 10.1101/2024.02.27.582233

**Authors:** Qiyao Wu, Jian Li, Tengfei Zhu, Huang Chang, Xu Wang, Jun Su

**Author notes:** **Correspondence:** Jun Su, Xu Wang. These authors have contributed equally to this work and share first authorship (Qiyao Wu, Jian Li, Tengfei Zhu, Huang Chang).

## Abstract

Plant undergo juvenile-to-adult transition to become competent for age-dependent floral induction and reproductive transition, which is of great significance for improving the seed quality and maintaining desirable genetic traits of Chinese fir, but the underlying molecular mechanize still remains unknown. Here, we investigated the function of our newly identified spermatophyte specific transcriptional co-activator TAC3 (Transcriptional Activator in Chinese fir 3) and its homologues (TAL1) in the model plant Arabidopsis. Both TAC3 and TAL1 can negatively regulate flowering, and activate miR156 expression to delay the phase transition. Moreover, we found that HDA9 and its its homologues in Chinese fir, ClHDA9, can directly binding to the promoter region of MIR156A and ClMIR156A, respectively. Directly interaction with ClHDA9 and HDA9 are necessary for the transcriptional activation of TAC3 and TAL1 on miR156, respectively. TAC3 and TAL1 also involve in the chromatin remodeling, shown as up-regulated H3K27ac level within the promoter region of ClMIR156A and MIR156A. Together, this work shows that TAC3 and its homologues are a new group of transcriptional co-activator that involving in aging-dependent flowering signal pathway of both angiosperms and gymnosperms.

## Introductions

Chinese fir (*Cunninghamia lanceolata* [Lamb.] Hook) is the second largest afforestation species in the world, accounting for approximately 24% of the total area of plantation forests in China. Its seed breeding is crucial for genetic breeding of forest trees in southern China^1, 2, 3^ . However, the maintenance of excellent traits in Chinese fir has long faced a technical bottleneck due to the low quality of the seeds^4^. Numerous studies have shown that asexual propagation commonly used in seed gardens, significantly shortens the nutrient growth cycle of Chinese fir. This leads to early flowering and incomplete nutrient growth, ultimately resulting in low seed quality^4, 5, 6, 7, 8, 9, 10^. Therefore, controlling the flowering time of Chinese fir is of great significance for improving the seed quality in seed gardens and maintaining desirable genetic traits during the breeding process.

Flowering time in plants is regulated by various of exogenous signaling pathways, including photoperiod, vernalization and gibberellin signaling pathways, as well as autonomous and age signaling pathways^11, 12, 13, 14^. Among these, the age signaling pathway is the most significant endogenous pathway and its molecular basis is also highly conserved in plants^6^. Vegetation growth can be divided into juvenile and adult stages, with the transition from juvenile to adult controlled by the age signaling pathway^15, 16, 17^. Chinese fir, like many perennial plants, requires approximately 20 years from seed germination for flower buds to begin forming. However, asexually propagated plants, mainly through grafting are capable of inheriting the age of the parent and start flowering in about 3-5 years^18, 19^. Therefore, the age signaling pathway plays a crucial role in the flowering process of Chinese fir. This hypothesis has also been widely confirmed in perennial woody plants such as poplar^20, 21, 22, 23, 24^.

The key component of the age signaling pathway, known as the miR156-SPL (SQUAMOSA PROMOTER BINDING-LIKE) cascade, has been found to be highly conserved in both annual and perennial plants^25, 26, 27, 28, 29^. During the juvenile stage, plants have high levels of miR156, which effectively represses the expression of its target SPLs (typically SPL9 and SPL15). As the plants enter adulthood, the expression of miR156 is significantly reduced, while the SPL level increases, thereby promoting flowering^16^. Additional, epigenetic regulation of miR156 has been observed in various plants, involving H3K27ac and H3K27me3. These modifications are responsible for promoting and repressing miR156 expression, respectively providing insights into the molecular mechanisms underlying the irreversible nature of age as a signaling factor.^29, 30, 31^. Several upstream transcription factors of miR156 have been identified^29, 32^, with most of them being transcriptional repressors such as *GCT* (*Arabidopsis MED12*) and *CCT* (*MED13*), as well as *Polycomb Repressive Complex 2* (PRC2). Additionally several epigenetic regulators, including HDA9, have been identified, which explains the down-regulation of miR156 expression with respect to plant age. However, the mechanisms behind the formation of high levels of miR156 are still unclear. Further investigation into the transcriptional activators of miR156 is necessary, especially in the context of perennial plants where our understanding is limited.^32, 33^.

In a previous study, we successfully identified a novel strong transcriptional activator, TAC3 (<u>T</u>ranscriptional <u>A</u>ctivator in <u>C</u>hinese fir <u>3</u>), in Chinese fir. In this study, we discovered that both TAC3 and its Arabidopsis homologue, TAL1 (<u>T</u>ranscriptional <u>A</u>ctivator of Chinese fir 3- <u>L</u>ike <u>1</u>), act as novel flowering repressors^34^. Additionally, we examined the age-related phenotypes of various transgenic materials and investigated the transcriptional regulation of ClmiR156 and AtmiR156 by TAC3 and TAL1, respectively. Through this analysis, we elucidated the underlying molecular mechanism of age signal-regulated flowering process in Chinese fir.

## Results

### TAC3 homologues TAL1 is a novel flowering time repressor in Arabidopsis

The function of TAC3 was explored through its homologues in the model plant Arabidopsis (TAL1)^34^. Firstly, the single mutant (*tal1*) and overexpression line (TAL1-OX) were developed (Fig. S1), and their flowering time phenotypes were measured under both long- and short-day conditions (Fig.1A & S2-3). TAL1-OX exhibited significantly longer flowering time than the wild type (WT) in both long- and short-day conditions, by 1.43 and 1.06 times, respectively; while *tal1* was 0.79 and 0.86 times significantly less than the WT, respectively (Fig.1B & S3B). Additionally, TAL1-OX had a higher number of rosette leaves than the wild type (WT) under both long- and short-day condition by 1.81 times and 1.20 times, respectively; while *tal1* was 0.76 and 0.83 times significantly less than the WT, respectively (Fig.1C & S3C). Furthermore, a transcriptome analysis was performed on *tal1* and WT by using RNA-seq, which revealed significant changes in aging signal (vegetation phase change) related genes in *tal1* (Fig. S4A).

**Fig. 1.**
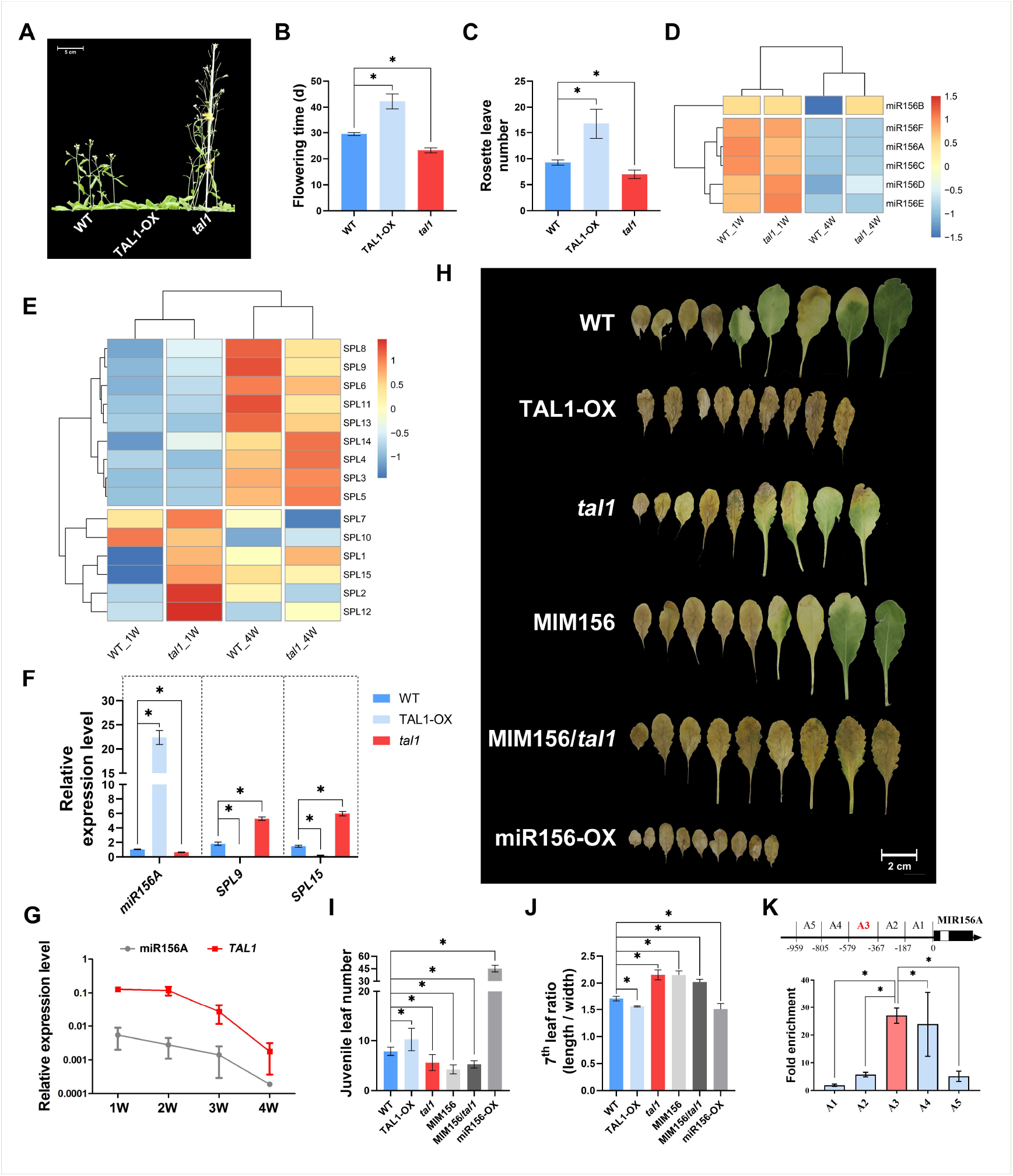
TAL1 suppresses flowering in Arabidopsis through the miR156-mediated aging signal pathway. The flowering phenotypes (A) of WT (wild type), TAL1 overexpression line (TAL1-OX), and TAL1 single mutation line (*tal1*) were compared based on their flowering time (B) and rosette leaf number (C). Both miRNA-seq and RNA-seq analyses were conducted using one-week-old WT (WT_1W) and *tal1* (*tal1*_1W), as well as four-weeks-old WT (WT_4W) and *tal1* (*tal1*_4W). The levels of different pri-miR156 (D) and SPL (E) levels were plotted using a heatmap. (F) The expression levels of aging signal response genes in different lines were also analyzed. (G) The dynamics of *miR156A* and *TAL1* level were quantified across different ages. Furthermore, the aging phenotypes of TAL1-OX, *tal1*, WT, MIM156 (mimic miR156), MIM156/ *tal1* (crossed line of MIM156 and *tal1*), and miR156-OX (miR156 overexpression line) were presented by first nine leaves in short day condition (H) and compered by their juvenile leaf number (I) and 7^th^ leaf ratio (J). (K) The ChIP-qPCR assay showed that TAL1 was able to bind to the promoter region of MIR156A. The ^*^ represents significant differences between samples at P < 0.05, based on a one-way ANOVA, with multiple comparisons made using Tukey’s test.

The expression patterns of mRNA and miRNA were compared between juvenile (one-week-old) and adult (four-weeks-old) plants. It was found that the levels of *miR156A* and miR156C (the major source of miR156) in *tal1* were significantly lower than in WT during the juvenile stage, but not during adulthood (Fig.1D). Additionally, most of the SPLs showed higher levels in *tal1* compared to WT during the juvenile stage (Fig.1E). The expression levels of *miR156A, SPL9*, and *SPL15* were measured in different genotypes. It was observed that in TAL1-OX, *miR156A, SPL9*, and *SPL15* expression level were 21.18, 0.01, and 0.04 times of that in WT, respectively. In *tal1*, these expression levels were 0.61, 0.35, and 0.25 times lower than in WT, respectively (Fig.1F). These results suggest that TAL1 may negatively regulate flowering through the miR156-SPL mediated aging signal.

To further investigate the hypothesis, the expression dynamic of *TAL1* regards to different age were quantified showed that *TAL1* level was decreased in an age-dependent manner (Fig.1G). Phenotypic analysis was performed and confirmed that TAL1-OX significantly delayed the phase transition, resulting in more juvenile leaves and lower 7^th^ leaf ratio compared to WT), while *tal1* could promote it in both long- and short-day condition (Fig.1H-J & S5-6). Interestingly, although TAL1 can interact with the promoter region (-579 ∼ -367 bp in the 5’ end of the cDNA) of MIR156A, no obvious difference in age phenotype was found between *tal1*, MIM156, or MIM156/*tal1*(Fig.1H-K & S5). Previous research has demonstrated that TAL1 functions as a transcriptional activator. Based on these findings, we conclude that TAL1 activates the expression of *MIR156A*, leading to a delay in the phase transition^34^.

### TAC3 activates miR156 to suppress the flowering time in Chinese fir

To investigate whether TAC3 has a similar function to TAL1, we conducted experiments using a heterologous expression system in Arabidopsis. Initially, we observed that TAC3 and TAL1 has similar subcellular localization, both in the cell membrane and nuclei (Fig. S7). It was found that, overexpressing TAC3 (TAC3-OX, Fig.S8) led to a significant delay in flowering and phase transition in short-day conditions compared to WT. This delay was characterized by a longer flowering time (1.07 times), an increased number of rosette leaves (1.24 times), more juvenile leaves (1.22 times), and a higher 7^th^ leaf ratio (0.89 times) compared to the WT (Fig. 2D-F & S9-10). Furthermore, TAC3-OX was able to rescue both the flowering and aging phenotypes of *tal1* in both long- and short-day conditions (Fig.1 A-F & S9/11). Interestingly, TAC3-OX/*tal1*exhibited similar levels of miR156, *SPL9*, and *SPL15* compared to the WT, while TAC3-OX shown significantly higher miR156 but lower *SPL9* and *SPL15* levels compared to the WT.

**Fig. 2.**
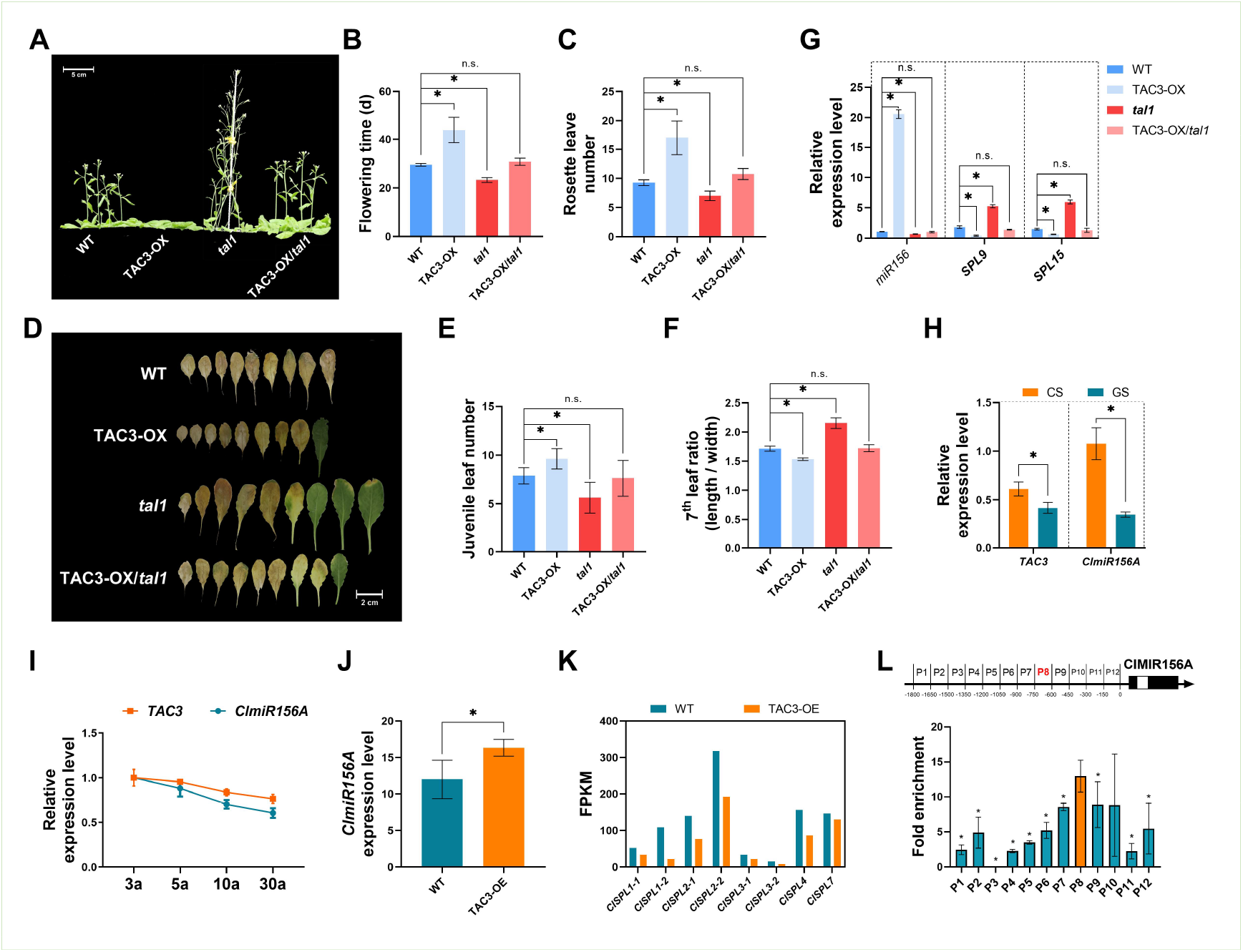
TAC3 positively regulated ClmiR156 level (miR156 of Chinese fir) in both the flowering and aging signal pathways. The function of TAC3 was investigated in Arabidopsis. The flowering phenotypes (A) of different lines including WT (wild type), TAC3 overexpression line (TAC3-OX), TAL1 single mutation line (*tal1*) and TAC3-OX/*tal1* (crossed line of TAC3-OX and *tal1*) were compared based on their flowering time (B) and rosette leaf number (C). Meanwhile, the aging phenotypes of TAC3-OX, *tal1*, TAC3-OX/ *tal1*, and WT were assessed by examining the first nine leaves in short day conditions (D) and comparing by their juvenile leaf number (E) and 7^th^ leaf ratio (F). (G)The expression levels of aging-related genes were analyzed in WT, TAC3 overexpression line, TAL1 single mutation line and TAC3-OX/*tal1*. Meanwhile, the levels of *TAC3* and ClmiR156A were quantified in GS and CS (H), as well as in samples of different ages (I). Transient expression assays were used to construct TAC3-OX Chinese fir seedlings, and the levels of ClmiR156A (J) and *ClSPLs* (K, quantified by RNA-seq and represented by FPKM value) were measured. (L) ChIP-qPCR assay revealed that TAC3 was able to bind to the promoter region of ClMIR156A. The ‘^*^’ and ‘n.s.’ represents significant differences between samples at P < 0.05 and P>0.05, respectively, based on a one-way ANOVA, with multiple comparisons made using Tukey’s test.

To further investigate the molecular mechanism of age-regulated flowering in Chinese fir, the overlapped differentially expressed miRNA between samples of different ages (3a and 30a) and different flowering times (grafting, GS, and cultivating, CS, seedlings) of Chinese fir materials was quantified (Fig. S12-13). The results showed that both TAC3 and ClmiR156 levels were higher in CS compared to GS and decreased along with the plant age (Fig.2H & I). Additionally, overexpressing TAC3 in Chinese fir resulted in 1.36 times higher *ClmiR156A* level compared to the WT, and most of the ClmiR156 targets (*ClSPLs*) were significantly down-regulated according to RNA-seq data (Fig. 2J-K & S14). In-vivo ChIP-qPCR assay results demonstrated that TAC3 could bind to the promoter region of ClMIR156A (-750 ∼ -600 bp in the 5’ end of the cDNA). Combining the strong transcriptional activation ability of TAC3, these findings suggest that TAC3 can activate ClmiR156 expression, leading to a delay in the flowering time of Chinese fir (Fig. 2L & S15).

### HDA9 bridges the transcriptional regulation of *MIR156A* by TAL1

Since both TAC3 and TAL1 lack a predictable DNA binding domain^34^, the question arises as to how they bind to the promoter region of *MIR156A*. By comparing the chromatin modification, it was observed that overexpression of TAL1 increased the H3K27ac level of the *MIR156A* promoter region while reducing the H3K27me3 level. In contrast, mutation of TAL1(*tal1*) showed the opposite pattern (Fig. S16). Thus, we screened the interacting protein of TAL1 within those reported proteins which could manipulate MIR156A expression by epigenetic regulation, and HDA9 was confirmed to directly interact with TAL1 by in-vitro assay (Fig.3A). To investigate whether HDA9 mediates the transcriptional regulation of miR156 by TAL1, a HDA9 deficient mutant (*hda9*) was developed by CRISPR-Cas9 and crossed with *tal1* (*tal1hda9*) or TAL1-OX (TAL1-OX/*hda9*) lines (Fig. S17). Mutation of HDA9 limits the function of TAL1 in both flowering and aging processes. This is evident from the similar flowering and aging phenotypes observed both in *tal1hda9* and TAL1-OX/*hda9* mutants, which closely resemble the wild type (WT) or *hda9* than *tal1* or TAL1-OX, respectively (Fig.3B-G & S18 -19). The significantly lower expression level of *miR156A* in TAL1-OX/*hda9* compared to TAL1-OX suggests that TAL1 is unable to activate *miR156A* expression in the presence of *hda9* (Fig. 3H). Additionally, the binding affinity of TAL1 to the promoter region of *MIR156A* is highly suppressed in *hda9* (Fig. 3I). Furthermore, the levels of H3K27ac and H3K27me3 in the binding region of TAL1 on the *MIR156A* promoter were measured, shown that H3K27ac levels in TAL1-OX/*hda9* was extremely lower but H3K27me3 were higher compared to TAL1-OX (Fig. S20). Taken together, the result indicates that HDA9 is necessary for TAL1 to bind to the promoter region of *MIR156A*.

**Fig. 3.**
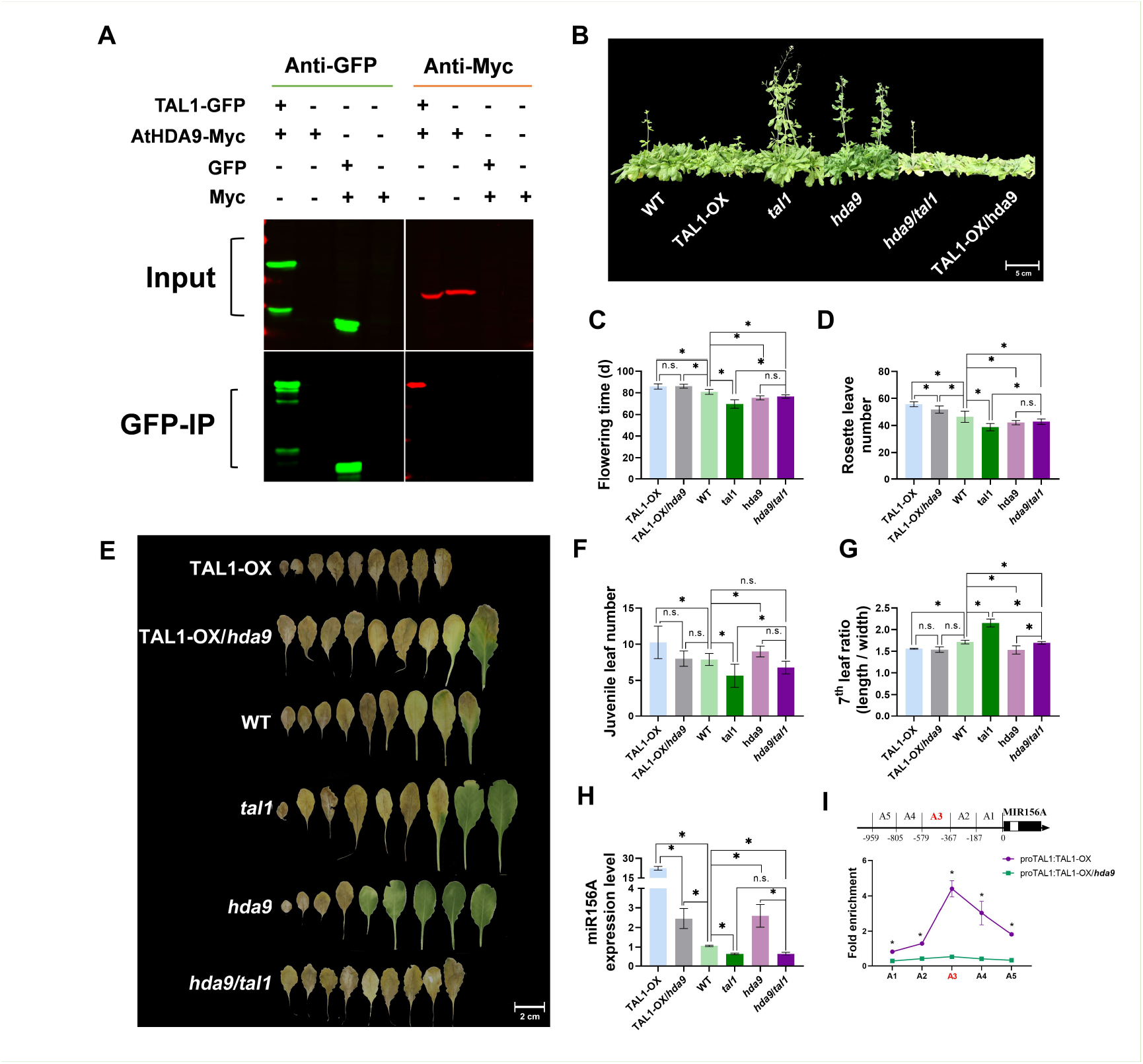
TAL1 binding to MIR156A promoter through HDA9 involvement in the aging signal pathway of Arabidopsis. (A) In-vitro protein-protein interaction assay showed that TAL1 can physically interact with Arabidopsis HDA9 (AtHDA9). The flowering (B) or aging phenotypes (E, presented by the first nine leaves in short day condition) of WT (wild type), TAL1 overexpression line (TAL1-OX), TAL1 single mutation line (*tal1*), HDA9 overexpression line (HDA9-OX), HDA9 single mutation line (*hda9*), and double mutation line (*hda9tal1*), TAL1-OX/*hda9* (crossed line of TAL1-OX and *hda9*), and HDA9-OX/*tal1* (crossed line of HDA9-OX and *tal1*) were compared in short day condition based on their flowering time (C), rosette leaf number (D), juvenile leaf number (F), 7^th^ leaf ratio (G), and expression level of miR156A (H). (I) The binding affinity of TAL1 on the promoter region of MIR156A was quantified by ChIP-qPCR with (TAL1-OX) or without (TAL1-OX/*hda9*) HDA9. The ‘^*^’ and ‘n.s.’ represents significant differences between samples at P < 0.05 and P>0.05, respectively, based on a one-way ANOVA, with multiple comparisons made using Tukey’s test.

### ClHDA9 involves in aging signal pathway and necessary for transcriptional regulation of *ClMIR156A* by TAC3 in Chinese fir

Similar to Arabidopsis, the level of H3K27ac and H3K27me3 in the TAC3 binding region on *ClMIR156A* (Chinese fir *MIR156A*), which vary with the age of Chinese fir, exhibit an inverse pattern. H3K27ac decreases with age, while H3K27me3 increases (Fig. S21A-B); Additionally, in-vivo overexpression of TAC3 (TAC3-OX) leads to increase the H3K27ac but decreased level of H3K27me3 level (Fig. S21C). To further investigate whether ClHDA9, the homologue of HDA9 in Chinese fir, has a similar function, we cloned ClHDA9 through phylogenetic analysis (Fig. S22). We confirmed that TAC3 can directly interact with ClHDA9 through an in-vitro protein-protein interaction assay (Fig.4A). Furthermore, ClHDA9 was overexpressed (ClHDA9-OX) in Arabidopsis *hda9* and successfully compensated for the flowering and age phenotypes of *hda9*. ClHDA9-OX/*hda9* exhibited similar flowering (flowering time and rosette leaf number) and age phenotypes (juvenile leaf number and 7^th^ leaf ratio) to the WT (Fig. 4B-D & S23-26). Also, the *miR156A* level in ClHDA9-OX/*hda9* is significantly lower than in *hda9* but similar to WT (Fig.4E). These results indicate that ClHDA9 plays a similar role in both the flowering and aging signaling pathways as HDA9.

**Fig. 4.**
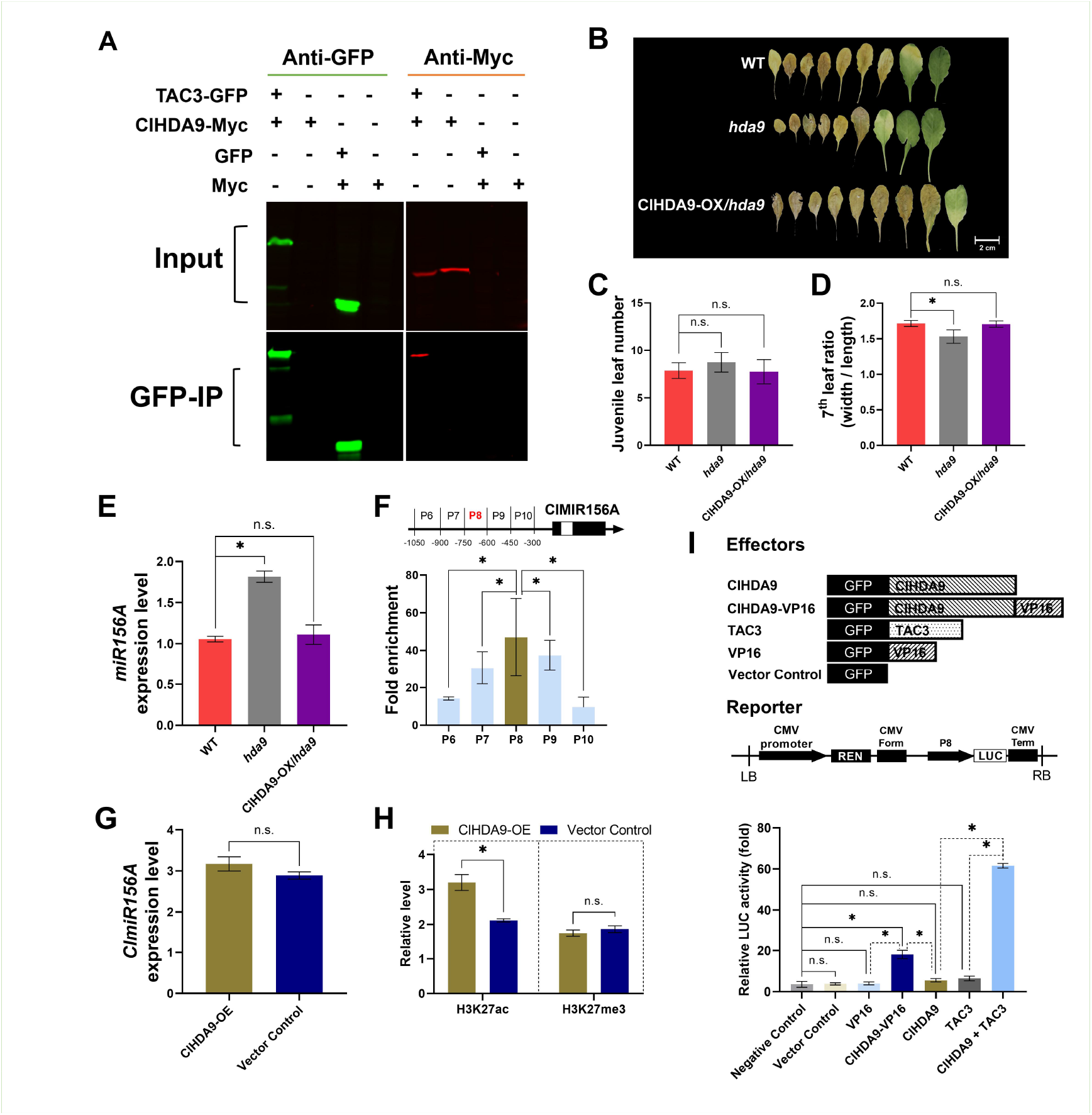
ClHDA9 is involved in the aging signal pathway and is necessary for the transcriptional regulation of ClMIR156A by TAC3 in Chinese fir. (A) In-vitro protein-protein interaction assay, it was shown that TAC3 can physically interact with ClHDA9 in Chinese fir. The aging phenotypes of WT (wild type), ClHDA9 overexpression line (HDA9-OX), HDA9 single mutation line (*hda9*), and ClHDA9-OX/*hda9* (the crossed line of ClHDA9-OX and *hda9*) were presented by the first nine leaves in short day conditions (B), and compared based on their juvenile leaf number (C), 7^th^ leaf ratio (D), and expression level of miR156A (E). (F) In-vivo ChIP-qPCR assay in the ClHDA9 overexpression Chinese fir line (ClHDA9-OX) showed that ClHDA9 can bind to the promoter region of ClMIR156A. The relative ClmiR156A level in WT and ClHDA9-OX Chinese fir line (G) was quantified. (H) The dynamics of H3K27ac and H3K27me3 levels in the promoter region of MIR156A were measured in Chinese fir by ChIP-qPCR. (I) In-vitro Dual-LUC assays shown that the transcriptional activation of ClmiR156A by TAC3 in Chinese fir. TAC3. proClmiR156A(P8-9) derived luciferase reporter gene (LUC) were co-expressed in 293T in various combinations with VP16, ClHDA9, ClHDA9-VP16 (transcriptional activation domian), and TAC3, to measure the LUC activity. And shown that Transcriptional activation of ClmiR156 by TAC3 requires ClHDA9. To normalize the activities of LUC, a constitutively expressed *Renilla* luciferase reporter gene (REN) was used as a reference (n = 9; mean ± SD). As controls, co-expression of GFP with the reporter served as the vector control, while the reporter alone was used as the negative control. The ‘^*^’ and ‘n.s.’ represents significant differences between samples at P < 0.05 and P>0.05, respectively, based on a one-way ANOVA, with multiple comparisons made using Tukey’s test.

In Chinese fir, ClHDA9 was overexpressed by in-vivo transient expression assay, to generate ClHDA9-OE. Then ChIP assay was conducted by using ClHDA9-OE, and shown that ClHDA9 shares the same binding site with TAC3 on the promoter region of *ClMIR156A* (Fig. 4F). Interestingly, we observed that ClHDA9-OE showed a significantly lower H3K27ac level in the ClHDA9 binding site of *ClMIR156A* compared to the vector control (Fig. 4H). However, we did not observe any noticeable differences in H3K27me3 or *ClmiR156A* levels between ClHDA9-OE and the vector control (Fig. 4G-H). To investigate the direct interaction between ClHDA9 and the promoter region of *ClMIR156A* (P8), we performed an in-vitro dual-LUC assay (Fig. 4I). We co-expressed ClHDA9 or its fusion protein with the universal transcriptional activator VP16 (ClHDA9-VP16) with the P8 driven *LUC* reporter. We found that VP16 or ClHDA9 alone did not activate LUC expression, but the fusion protein ClHDA9-VP16 significantly induced the LUC reporter level, showing a 4.47-fold and 3.24-fold increase compared to VP16 and ClHDA9, respectively. Furthermore, in the same system, TAC3 alone did not induce LUC expression, but co-expression of TAC3 and ClHDA9 increased the LUC level, showing a 9.47-fold and 11.02-fold increase compared to TAC3 and ClHDA9 alone, respectively. This study demonstrates that ClHDA9 can interact with TAC3 and directly bind to the promoter region of *ClMIR156A*, indicating its essential role in the transcriptional activation of TAC3 on ClMIR156A (Fig. 5).

**Fig. 5.**
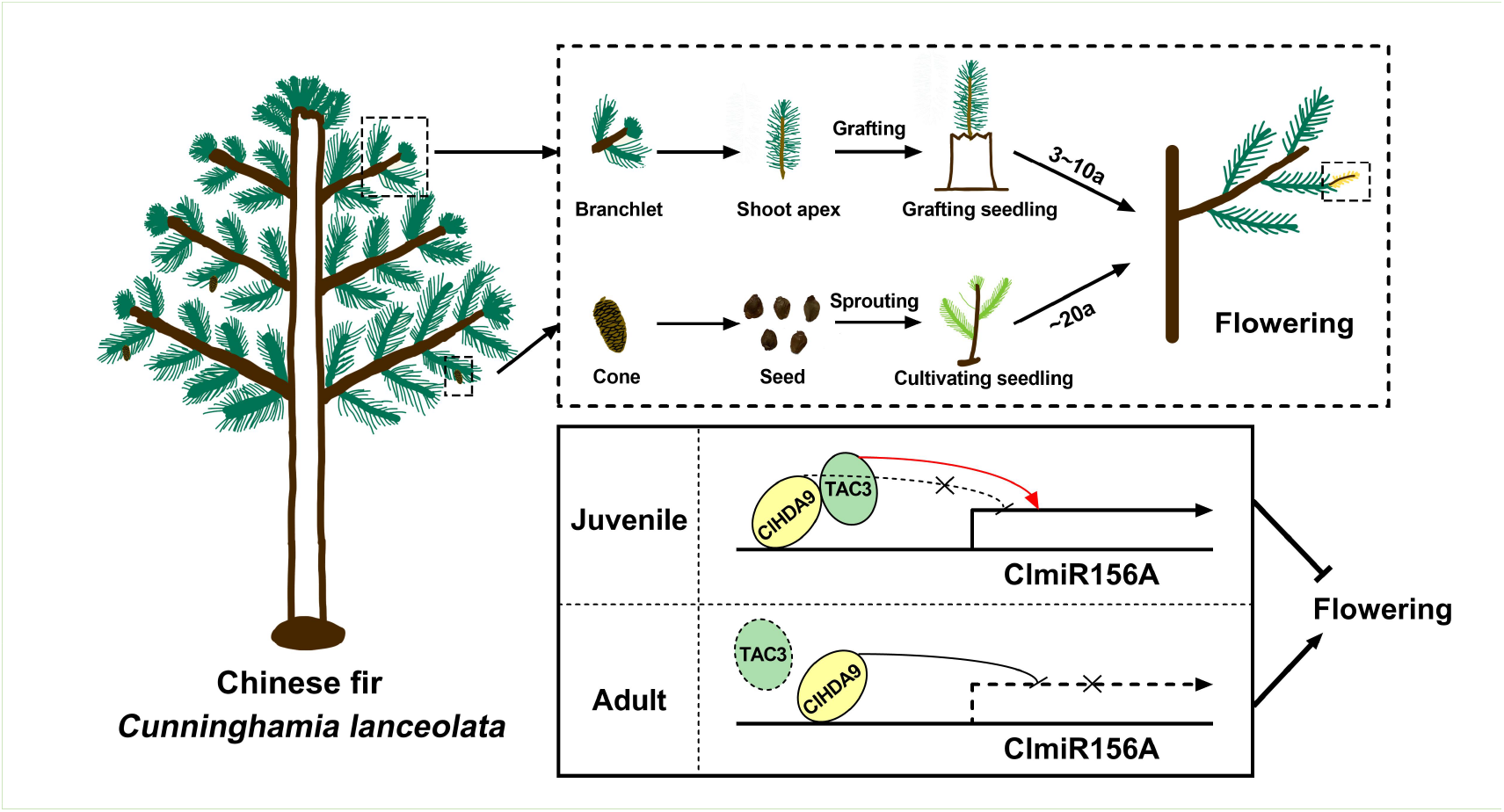
Illumination of the TAC3-based aging signal pathway regulates the flowering time of Chinese fir.

## Discussion

Grafting is a commonly used asexual propagation technique for woody plants. However, it can result in early flowering, leading to low seed quality. This limitation significantly hinders the maintenance of good genetic traits in traditional tree breeding. One possible explanation is that grafting tissues may inherit the “age” of the stock plant causing the grafted plant to flower earlier. However, the underlying molecular mechanize behind this phenomenon still remain a myster^35, 36^. In this study, we investigated the function of TAC3 in Chinese fir and its homologues (TAL1/2) in the model plant Arabidopsis. We found that these genes act as flowering repressors by integrating the aging signal in plants (Fig. 1-2). MADS-box family proteins have previously been reported as flowering time regulators in Chinese fir, which is a typical flowering regulator in woody plants^21, 37^. Additionally, MADS-box proteins have also been implicated in the aging process of several woody plants^21, 38^. TAC3 and its homologues are atypical regulators of flowering time, and they are highly conserved in plants^11, 12, 34^. Therefore, there is an urgent need for further exploration of their additional functions. Transcriptome data suggests that the photoperiod signal is involved in TAL-regulated flowering time control (Fig. S4B). However, no significant difference in flowering time phenotypes was observed between TAL1-OX or *tal1* under long- and short-day conditions (Fig.1 & S3). Apart from the aging process, cold accumulation is another important factor for flowering in Chinese fir. Our data indicates that TALs may be involved in the vernalization signal pathway (Fig. S4B)^18, 39^.

The most promising aging phenotype in model plants is based on the first abaxial leaf trichome, which represents the vegetative phase transition (from the juvenile to adult stage)^16, 40^. However, the aging phenotype is different in woody plants, making it difficult to clearly identify the phase transition in Chinese fir^41, 42^. Since the function of miR156 in aging signal is highly conserved across different plant species, we use the ClmiR156 level as the positive control or internal gene marker in the Chinese fir system to study the aging signal, especially in Chinese fir plants of different ages (Fig. 2, 4, S12 - 21)^43^. To investigate the environmental cues of plant endogenies aging signal, many transcriptional regulators of miR156 have been documented. However, most of them directly interact with the DNA of the MIR156s promoter region and response to specific environmental cues that affect the aging process^29, 31^. TAC3 and TALs are the first reported transcriptional co-activator of miR156. They appear to integrate multiple environmental cues to regulate the aging signal by interacting with various DNA binding proteins (Fig. S4B). Interactome and R-ChIP are required to screen TAC3 and TALs associated DNA binding proteins of different MIR156 promoters, to further uncover the crosstalk of different environmental cues affect the aging process and its underline molecular mechanisms^44^.

In this study, we developed a deficient mutant of Arabidopsis HDA9 (*hda9*), with a single amino acid substitution, HDA9^Q337H^. Similar to other HDA9 mutants, *hda9* exhibits an early flowering phenotype in short-day conditions, but a delayed phase transition (Fig.3B-G & S25 -26) ^30, 45^. This suggests that the 337th amino acid, glutamine, is crucial for the function of HDA9 in both aging and flowering signal pathways. Interestingly, HDA9^Q337H^ displays an early flowering phenotype in long-day condition, while other reported *hda9* mutants show a similar flowering phenotype to the WT (Fig. S27). This further highlights the importance of the 337th amino acid glutamine in HDA9’s response to photoperiod^45^.

The flowering and aging phenotypes of TAL1-OX/*hda9* were similar to *hda9*, while the aging phenotype of *tal1hda9* was similar to *tal1*, indicates TAL1 functions in both a HDA9-dependent and independent manner (Fig.3). Previous studies have reported that the HDA9 binding site of MIR156A/C is located within the gene body. However, our work demonstrates that TAL1 can interact with HDA9 within the 5’ up-stream of the MIR156A gene body to regulate miR156A expression and chromatin remodeling (Fig. 3I & S16)^30^. Histone deacetylase family proteins exhibit high functional diversity and can function as both chromatin deacetylases and acetylases for the same member^46, 47^. In this study, HDA9 was found to reduce the H3K27ac level of the MIR156A promoter in Arabidopsis, while its homologues in Chinese fir (ClHDA9) increased the H3K27ac level of the ClMIR156A promoter (Fig. 4H & S8).

The lack of a stable in-vivo gene transformation system in Chinese fir significantly limits our methodology for exploring TAC3 function. In this study, we investigated the function of TAL1 in flowering and aging using transgenic Arabidopsis materials (Fig. 1 & 3). Furthermore, we studied the function of TAC3 flowering and aging by overexpressing it in related Arabidopsis transgenic lines. We also examined the TAC3 expression pattern of TAC3 and the epigenetic regulation of ClmiR156 in different Chinese fir materials (Fig. 2, 4, S21C). One challenge with this approach is that we cannot obtain homozygous seeds for cultivating plants in Chinese fir due to self-incompatibility. However, this issue could potentially be resolved by developing a seed orchard of Chinese fir using cultivars that have similar flowering times^18, 48, 49, 50^.

## Materials and methods

### Plant material and growth conditions

All Arabidopsis stocks used in this study were on the WT (wild type, Col) back ground. The MIM156 (*mimic miR156*) and miR156-OX (*miR156* overexpression line) have been previously described^51^. The TAL1 single mutation line (*tal1-1*, SALK_052626) was obtained from the ABRC (https://abrc.osu.edu/). Fragments of the *AtTAL1* (AT1G13360), *TAC3*, and *ClHDA9* coding regions were cloned and transformed into WT to generate transgenic lines with TAL1 overexpression (TAL1-OX), TAC3 overexpression (TAC3-OX), and ClHDA9 overexpression (ClHDA9-OX) by the floral dip method. The HDA9 single mutation line (*hda9*) and double (*hda9/tal1*) mutation line were constructed in WT and *tal1* backgrounds using the *pYAO*-based CRISPR/Cas9 and floral dip method, respectively. MIM156/*tal1* (crossed line of MIM156 and *tal1*), TAC3-OX/*tal1* (crossed line of TAC3-OX and *tal1*), HDA9-OX/*tal1* (crossed line of HDA9-OX and *tal1*), TAL1-OX/*hda9* (crossed line of TAL1-OX and *hda9*), and ClHDA9-OX/*hda9* (crossed line of ClHDA9-OX and *hda9*) were generated through the crossing method. Seeds were sown on PINDSTRUP soil, stratified at 4°C for 3 d, and then transferred to long-day (LD, 16 h light–8 h dark) or short-day (SD, 8 h light–16 h dark) growth conditions at 22°C.

Chinese fir seeds were collected from the Chinese fir gene bank in Guanzhuang state-owned forest farm in Shaxian, Fujian, China (117° 35′∼117° 50′ E, 26° 24′∼27° 38′ N). The gene bank established in 1970 by the Guanzhuang state-owned forest farm and Fujian Academy of Forestry Sciences, is located in a subtropical maritime monsoon climate. The Chinese fir seeds were sown on PINDSTRUP soil, stratified in darkness for 1 day, and then transferred to long-day (LD, 16 h light–8 h dark) growth conditions at 22 °C. As well as, Chinese fir seedlings of different ages (1a, 2a, 3a, 5a, 10a, 30a) obtained from this gene bank, including grafting (GS) and cultivated (CS) seedlings were collected for qRT-PCR and chromatin IP (ChIP) analysis. The GS was grafted with newly developed branches from 10-year-old Chinese fir plants of the same cultivar as the CS and grown in the same year.

### Plasmid construction and transformation

The full-length coding regions of *AtTAL1* (585 bp), *AtHDA9* (1281 bp), *TAC3* (537 bp), and *ClHDA9* (1296 bp) were PCR-amplified from cDNA of Arabidopsis WT plants and Chinese fir seedlings using 2×Profusion™ PCR Master Mix (HRF0088, Herui, Shanghai, China), respectively. The *promiR156A* (450-750 bp fragment upstream of the transcription start site) was PCR-amplified from DNA of Chinese fir seedlings. The fragment of VP16 (234 bp) was descrbed previously^52^. The full-length coding regions of *TAL1, TAC3, AtHDA9*, or *ClHDA9* can be found in Supplementary Table.S1.

To generate *35S::TAL1-GFP* and *35S::TAC3-GFP*, the full-length coding regions of *TAL1* and *TAC3* ligated into SpeI- and SmaI-digested *pFGFP* (R3133, R0141, New England BioLabs, Ltd., USA) using the In-Fusion method. To generate *ACT2::TAL1-4MYC* and *ACT2::ClHDA9-4MYC*, the full-length coding regions of *TAL1* and *ClHDA9* ligated into SmaI- and BamHI-HF-digested *pDT1H-4MYC* (R0141, R3136, New England BioLabs, Ltd., USA) using the In-Fusion method^53^. Similarly, the full-length coding regions of *AtHDA9* and *ClHDA9* ligated into SpeI-digested *pCMV-4MYC* to produce the *CMV::AtHDA9-4MYC, CMV::ClHDA9-4MYC* vector. The full-length coding region of *TAL1* and TAC3 ligated into SpeI-digested *pQCMV-GFP* to produce the *QCMV::TAL1-GFP* and *QCMV::TAC3-GFP* vector. Additionally, the fragment of *VP16*, the fusion fragment of the promoter region of *miR156A* and *VP16* ligated into SpeI-digested *pQCMV-GFP* to produce the *QCMV::GFP-VP16* and *QCMV::GFP-ClHDA9-VP16* binary vector. The promoter region of *ClMIR156A* ligated into SpeI-digested *pQCMV-LUC* to produce the *QCMV::promiR156A-LUC* vector. All the primers used in the study are shown in Supplementary Table.S2.

### Phenotypic investigation

In the analysis of Arabidopsis flowering phenotypes, plant age was measured from the time of transfer to the growth chambers after sowing. Once the main inflorescence became visible approximately 8-10 cm, the flowering time was recorded. Subsequently, the rosette leaves were detached, fully expanded, and attached to cardboard using double-sided tape. These leaves were then photographed and counted. Furthermore, the flowering phenotypes of all lines were continuously recorded and photographed for each line from the time of sowing.

In the analysis of Arabidopsis age phenotypes, the number of juvenile leaves, or the leaves on which the cotyledon trichomes first appeared were counted. The length and width of the fourth leaf were measured using ImageJ software (version 64-bit Java8), and the leaf aspect ratio was calculated by averaging these measurements. Three biological replicates of each genotype were analyzed, with a total of 30 biological replications used for each strain.

### In-vivo transient expression assay

To transform the plasmids (*TAC3-GFP* and *pFGFP*), 10 μg of each were added to 10-day-old clean seedlings of Chinese fir using Zhu’s method with minor modifications^52^. The polypeptide and plasmids were added to a 5ml falcon tube containing 300 μl of 1×PBS buffer, with a ratio of 1:1. The falcon tube was then transferred to 28°C for 30 minutes. Then the tube was transferred to room temperature and the volume was increased to 5 ml using ddH_2_O. Vacuum treatment was performed for 5 minutes. Following incubation in the dark for two days, Western blot analysis was conducted to detect the GFP cassettes in the seedlings and confirm the successful transformation.

### Quantitative real-time PCR

Leaves from grown Arabidopsis plants, branches of different ages (1a, 2a, 3a, 5a, 10a, 30a), as well as grafting (GS) and cultivating (CS) seedlings of Chinese fir were collected. Total RNA was extracted using the FastPure Plant DNA Isolation Mini Kit (DC104-01, Vazyme, Nanjing, China) according to the manufacturer’s protocol. For cDNA synthesis, 1 μg of total RNA was used with the cDNA Synthesis SuperMix for qPCR Kit (11141ES, Yeasen, Shanghai, China). Quantitative real-time PCR (qRT-PCR) was performed using the QuantStudio 6 system and SYBR-Green PCR Mastermix (11202ES08, Yeasen, Shanghai, China), with *GAPDH* and *ClGAPDH* serving as the reference genes for Arabidopsis and Chinese fir, respectively^54, 55^. Three technical replications were conducted. The relative expression level or fold change was calculated using the comparative CT method (2^−ΔΔ*CT*^)^56^. The sequences of primers for qRT-PCR are listed in Supplementary Table.S2.

### RNA-seq and analysis

Arabidopsis leaves from 1-week and 4-week-old plant lines (WT and *tal1*) in SD conditions, two-week-old treated and untreated Chinese fir seedlings (TAC3-OX, WT) were harvested and crushed in liquid nitrogen for further sequencing. Total RNA extraction was used according to procedures described previously in ‘Quantitative RT-PCR’

The degradation and contamination of RNA were monitored on 1% agarose gels and the quality and integrity of RNA were assessed using a NanoDrop spectrophotometer (840-317400, Thermo Scientific, DE, USA). A total amount of 1.5 μg RNA per sample was used as input material for the RNA sample preparations. Sequencing libraries were generated using the NEBNext® Ultra™ RNA Library Prep Kit for Illumina® (E7770, New England BioLabs, Ltd., USA) by the Beijing Allwegene Technology Company Limited (Beijing, China). The quality of the constructed libraries was assessed using the Agilent Bioanalyzer 2100 system and fastq files (raw data) were sequenced using Illumina Novaseq 6000 using the PE 150 sequencing strategy by the Allwegene Technology Company. Three technical replications were conducted for each RNA-seq library. Using perl, cutadapt, and perl scripts, raw data were cleaned up to obtained clean data for downstream analysis. The RNA sequencing files have been made available by the National Center for Biotechnology In·formation (NCBI) under accession number PRJNA1007949 (https://www.ncbi.nlm.nih.gov/). However, no publicly available genomic information is available, the RNA-seq analysis of Chinese fir used reference-free assembly and annotation.

For downstream analysis in Arabidopsis, clean reads were mapped to the Arabidopsis reference genome assembly and annotation (TAIR10) using STAR (v2.5.2b). The quantification of transcript abundance (readcount as well as FPKM) was conducted with HTSeq (v 0.5.4), and Differentially expressed genes (DEGs) analysis was performed using the DESeq2 package (R studio, https://bioconductor.org/packages/release/bioc/html/DESeq2.html). P<0.05 and |log2foldchange| ≥ 2 were set as the threshold for significantly differential expression. The clusterProfiler and the org.at.tair.db package (R stuido) for Gene Ontology (GO) enrichment analysis of the resulting differential genes.

For downstream analysis in Chinese fir, after the adapter sequences and low-quality sequences of raw reads were deleted, clean reads were assembled using Trinity (v2.8.5)^57^. CD-hit was used to classify transcripts, corset was used to remove redundancy, and BUSCO (Benchmarking Universal Single Copy Orthologs, V 3.0.2) was used for the evaluation of the quality of the constructed transcriptome sequences^58, 59^. Then, these sequences were annotated by commonly used databases, including NCBI protein non-redundant database (NR), and Gene Ontology (GO) (http://www.geneontology.org/). The transcripts obtained from Trinity assemble were used as reference sequences, and the clean reads were mapped to the reference sequences by RSEM (v1.2.15) to further count the number of readcounts for each gene in each sample, and the FPKM conversion was performed to analyze the expression level of the genes. DEGs analysis was performed using DESeq (1.10.1). GO enrichment analysis of the DEGs was implemented by the GOseq R packages^60^.

### miRNA-seq and analysis

Arabidopsis leaves from 1-week and 4-week-old plant lines (WT and *tal1*) in SD conditions, different age (3a, 30a) and grown conditions (CS, GS) Chinese fir seedlings were harvested and crushed in liquid nitrogen for further sequencing. Total RNA extraction was used according to procedures described previously in ‘Quantitative RT-PCR’ and RNA quality examining was used according to procedures described previously in ‘RNA-seq and analysis’. A total amount of 3 μg total RNA per sample was used as input material for the small RNA library by the Allwegene Technology Company. Sequencing libraries were generated using the NEBNext® Multiplex Small RNA Library Prep Set for Illumina (E7330S, New England BioLabs, Ltd., USA) following manufacturer’s recommendations and index codes were added to attribute sequences to each sample. Then library quality was assessed on the Agilent Bioanalyzer 2100 system using DNA High Sensitivity Chips. The library preparations were sequenced on an Illumina Novaseq 6000 platform by the Allwegene Technology Company and 50 bp single-end reads were generated.

For Arabidopsis, the small RNA tags were mapped to reference sequence by Bowtie (bowtie-0.12.9) without mismatch to analyze their expression and distribution on the reference genomic (TAIR10)^61^. Mapped small RNA tags were used to looking for known miRNA. miRBase20.0 was used as reference, modified software mirdeep2 (mirdeep2_0_0_5) and sRNA-tools-cli were used to obtain the details of potential miRNA^62^. For Chinese fir, the details of the sRNAs were obtained by using the assembled sequences as reference sequences for map annotation. In addition, miREvo and mirdeep2 software were used to predict the novel miRNAs in the annotated samples, and the sequence, length, and number of occurrences of the matched sRNAs were counted in each sample. The miRNA expression levels were estimated by TPM (transcript per million) through the following criteria: Normalization formula: Normalized expression = mapped readcount / Total reads^*^1000000^63^. The miRNA expression levels were counted using TPM (transcript per million), and differential expression analysis was performed using the DESeq R package (1.8.3), followed by GO enrichment analysis predicted by the target gene candidates of differentially expressed miRNAs

### In-vitro protein protein interaction assay

#### Cell culture and transfection

The HEK 293T cells, obtained from Dr. Qin Wang, were incubated in a CO_2_ incubator at 37°C in DMEM (CM-0001, Pricella, Wuhan, China). Prior to transfection, the 293T cells were cultured in 6 cm dishes (approximately 1.7×10^6^ cells) for 36 hours. Each dish received 5 μg of plasmid. For transfection, PEI MW40000 (40816ES02, Yeasen, Shanghai, China) was used as the transfection reagent at a ratio of 1:3 (Plasmids:PEI MW40000). After 24 hours of transfection, the cells were harvested and washed twice with 1×PBS buffer. The cell precipitate obtained after centrifugation, was used for subsequent Co-immunoprecipitation analysis.

### Co-immunoprecipitation (Co-IP)

The cells obtained from 293T transfection were collected for Co-IP analysis. 350 μl of 1%Brij buffer+inhibitors (1% Brij, 50 mM Tris, pH8.0, 150 mM NaCl, 1 mM EDTA and 1×PIC) was added to the microtube containing the cells and lysed for 30 min at 4°C. The crude extract was then centrifuged twice at 12 000 rpm for 10 min. After centrifugation, 30 μl supernatant of the supernatant was saved for Input. The remaining supernatant was collected for IP assays. The beads were washed three times with 1% Brij buffer (1% Brij, 50 mM Tris, pH8.0, 150 mM NaCl and 1 mM EDTA). Then, 20 μl of anti-GFP beads (AE074, ABclonal, Wuhan, China) were added to each tube for IP and gently rocked for 3h at 4°C. The beads were then washed five times with 1%Brij buffer and eluted with 40 μl of 1× Flag-peptide buffer (50× Flag-peptide and 1% Brij buffer+inhibitors, F3290, sigma, Danvers, MA, USA) rotating for 30 min at 25°C. The eluted substances, as well as inputs, were subjected to western blot analysis. Antibodies against GFP-Rabbit (AE011, ABclonal, Wuhan, China) and myc-mouse(AE010, ABclonal, Wuhan, China) were used as primary antibodies in western blot analysis. Subsequently, Goat Anti-Rabbit IgG(H+L) Alexa Fluor 680(33218ES, Yeasen, Shanghai, China) and Goat Anti-Mouse IgG(H+L) Alexa Fluor 790(33118ES, Yeasen, Shanghai, China) were used as secondary anti-bodies. Auto-fluorescence was detected and analyzed using the Odyssey® DLx Imaging System (LI-COR, Lincoln, Nebraska, USA) and Image Studio Lite 3.1 software.

### Chromatin immunoprecipitation–qPCR (ChIP-qPCR)

Here, 2–3 g 1-, 2-, 3-,and 4-week-old Arabidopsis leaves, two-week-old treated and untreated Chinese fir seedlings (TAC3-OX, WT) were harvested and crushed in liquid nitrogen. Then sample tissues were suspended in Nuclear Isolation and X-linking buffer (1 M sucrose, 10 mM Hepes pH 8.0, 5 mM KCl, 5 mM MgCl_2_, 1 mM PMSF, 1×PIC and 0.1% Triton X-100). Stop X-Linking with addition of 2M glycine. Pellets were washed with Extraction buffer 2+inhibitors (0.25 M sucrose, 10 mM Tris–HCl, pH 8.0, 10 mM MgCl2, 5 mM β-Mercaptoethanol, 1 mM PMSF, 1×PIC and 1% Triton X-100), and resuspended in nuclei lysis buffer (50 mM Tris–HCl, pH 8.0, 10 mM EDTA, 1% SDS, 1mM PMSF and 1×PIC). Then added dilution buffer (1.2 mM EDTA, 16.7 mM Tris–HCl, pH 8.0, 16.7 mM NaCl, 1.1% Triton X-100, 1mM PMSF and 1×PIC) to a final volume of 6ml and sonicated using a Covaris ultrasonicator M220. Save 5% supernatant for Input. Next, 1% of antibodies against GFP (AE079, ABclonal, Wuhan, China) was used in IP. The IP protein and DNA were reverse crosslinked, and DNA including IP and Input was isolated using the QIAquick PCR Purification Kit (28104, QIAGEN, Germany). qPCR was performed on QuantStudio 6 real-time system after DNA was extracted from the IP product. For GFP ChIP, the ChIP assay of pFGFP control vector was used as a negative control. *GAPDH* and *ClHDA9* was used as a control locus and data are presented as the ratio of (Target-GFP ChIP gene of interest/*pFGFP* ChIP gene of interest) normalized to the promoter locus^54, 55^. Primers for ChIP analysis are listed in Table.S2

### In-vitro Dual-Luciferase Reporter assay

The dual-luciferase assay was performed by introducing the constructs into HEK 293T cells. After transfection, the LUC and REN values were measured 24 hours later using the Dual-Luciferase® 80 Reporter Assay System (E1910, Promega, WI, USA)^34, 64^.

