## Supplemental Figures S1-S26 for "Novel transcriptional activator TAC3 regulates age-dependent floral transition in Chinese fir (*Cunninghamia lanceolata*)"

### Figure S1

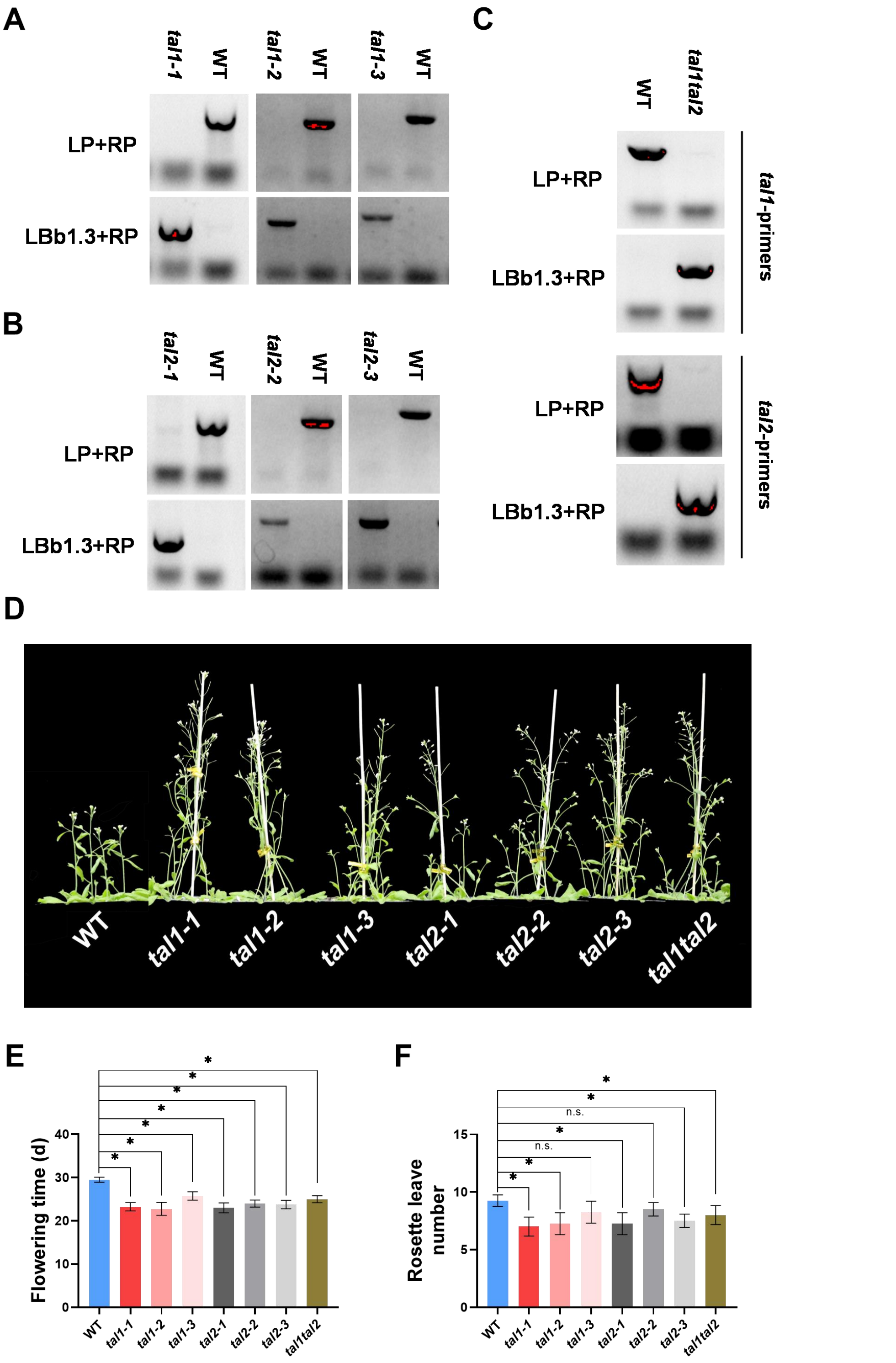


**Fig. S1 TALs suppress flowering in Arabidopsis grown at LD conditions.** (A), (B) and (C) Schematic diagram of *tal1-1*, *tal2-3* and *tal1tal2* pure haplogroup validation with LP + RP amplification of *TAL1* and *TAL2* genomic DNA and LBb1.3 + RP amplification of a small sequence of T-DNA insertion. The flowering phenotypes (D) of WT (wild type), TAL1 single mutation line (*tal1-1*, *tal1-2*, *tal1-3*), TAL2 single mutation line (*tal2-1*, *tal2-2*, *tal2-3*), and TAL1TAL2 double mutation line (*tal1tal2*) were compered by their flowering time (E), rosette leave number (F). The '*' and 'n.s.' represents significant differences between samples at P < 0.05 and P>0.05, respectively, based on a one-way ANOVA, with multiple comparisons made using Tukey's test.

### Figure S2

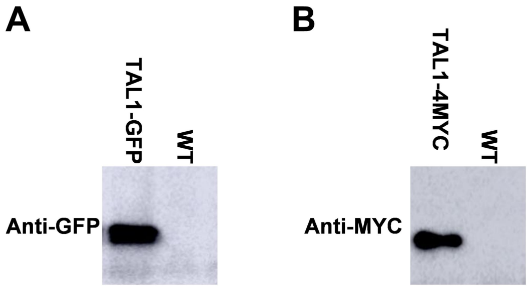


**Fig. S2 Identification of TAL1 overexpression line.** TAL1-GFP overexpression line (A) and TAL1-4MYC (B) were identified by Western Blot.

### Figure S3

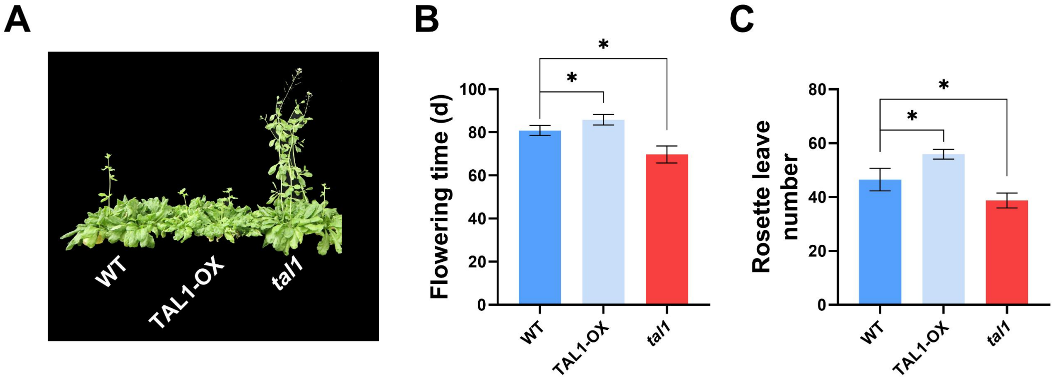


**Fig. S3 TAL suppress flowering in Arabidopsis at SD conditions.** The flowering phenotypes (A) of WT (wild type), TAL1 overexpression line (TAL1-OX) and TAL1 single mutation line (*tal1*) were compered by their flowering time (B), rosette leave number (C). The '*' represents significant differences between samples at P < 0.05, based on a one-way ANOVA, with multiple comparisons made using Tukey's test.

### Figure S4

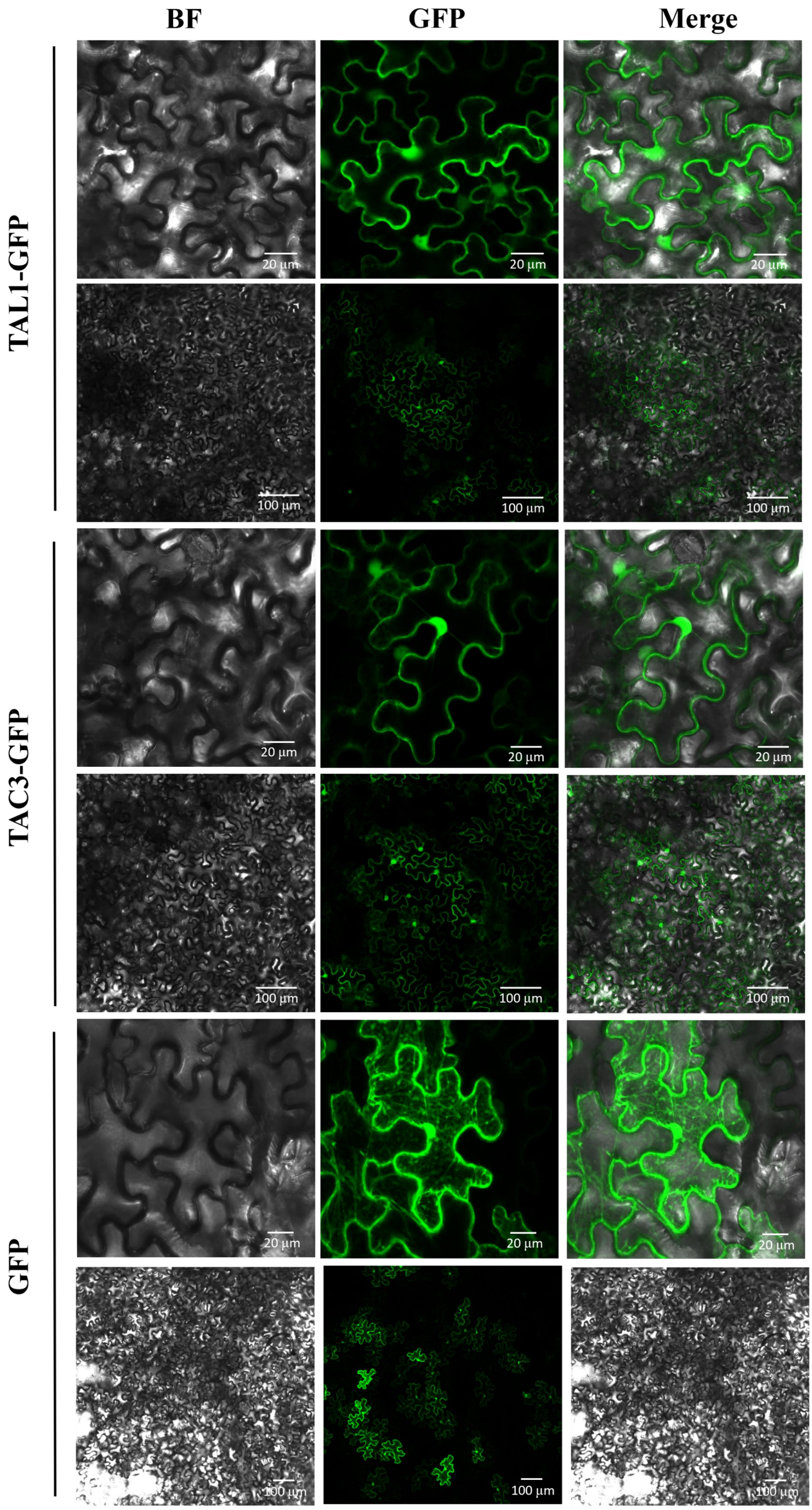


**Fig. S4 Subcellular localization of *TAL1-GFP*, *TAC3-GFP* and *pFGFP* in *Nictiana benthamiana* cells.**

### Figure S5

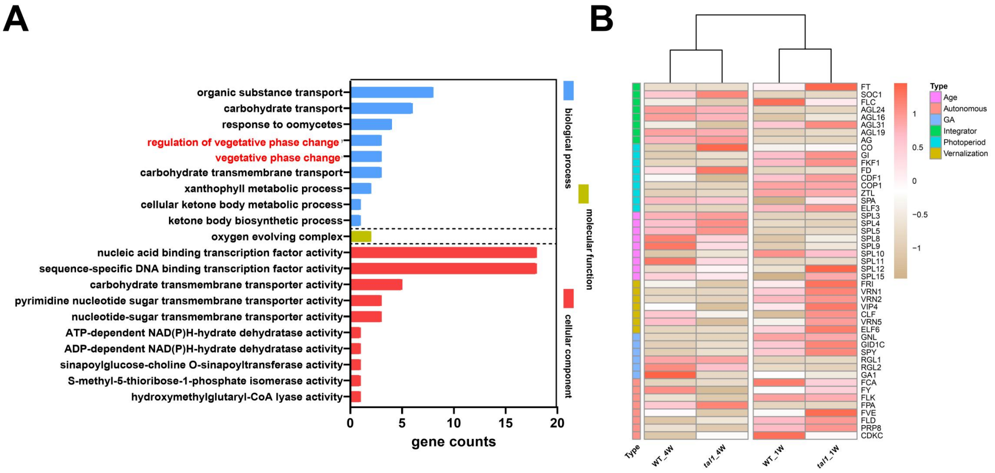


**Fig. S5 TAL1 suppress the flowering in Arabidopsis through vegetative phase change signal pathway and aging signal pathway by transcriptional analysis.** (A) The functions of differentially expressed genes between WT and *tal1* were predicted by GO enrichment analysis. The different color represents three categories: biological process, cell component, molecular function. The height of the column indicates the number of differentially expressed genes in the pathway. (B) Heatmap showing the expression profiles of selected flowering time related genes through different pathway in *tal1* and WT plants. The key below shows down-regulation to the left and up-regulation to the right.

### Figure S6

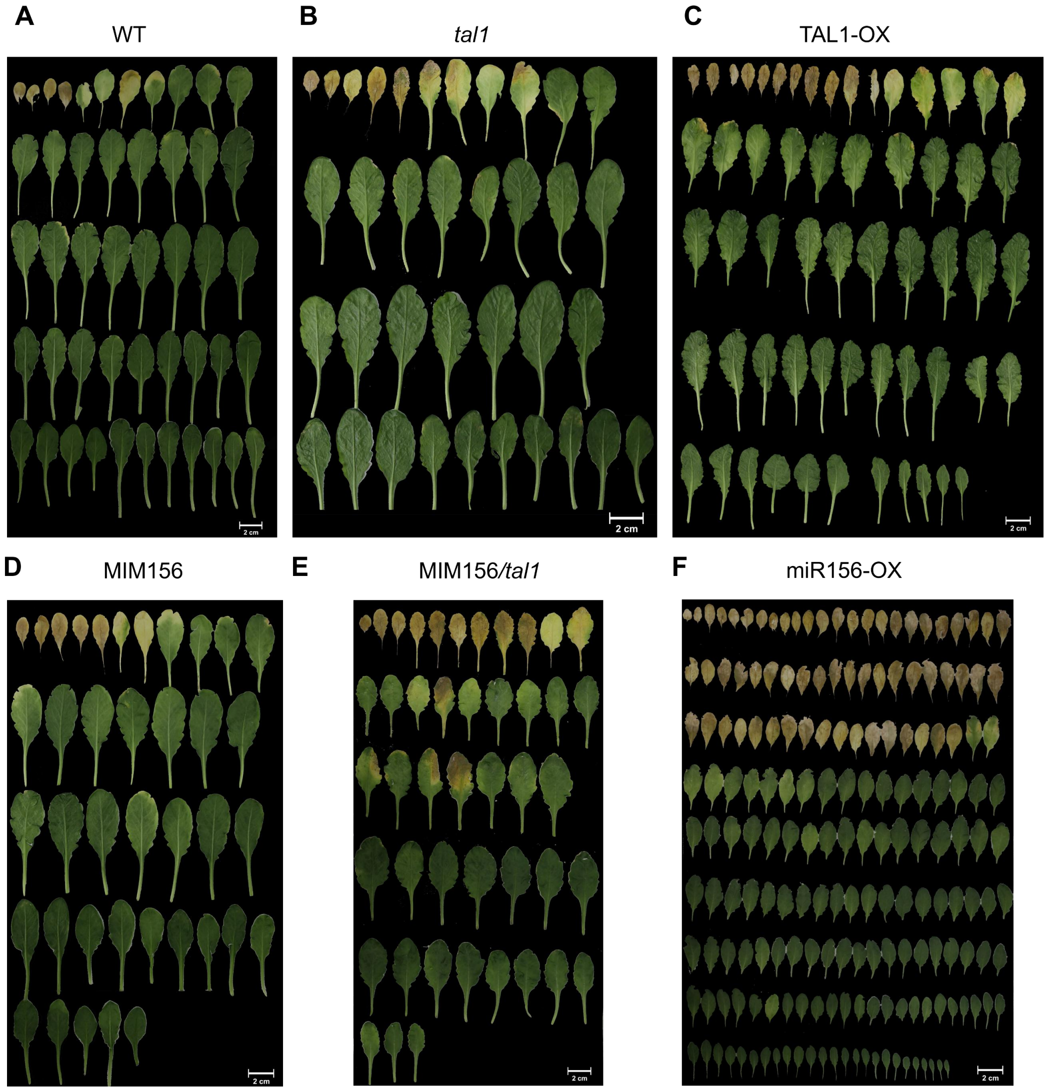


**Fig. S6 The aging phenotypes of TAL1-related lines in Arabidopsis at SD conditions**. The aging phenotypes of WT (A), *tal1* (B), TAL1-OX (C), MIM156 (mimic miR156) (D), MIM156/ *tal1* (crossed line of MIM156 and *tal1*) (E), and miR156-OX (miR156 overexpression line) (F) were presented by all leaves in short day condition.

### Figure S7

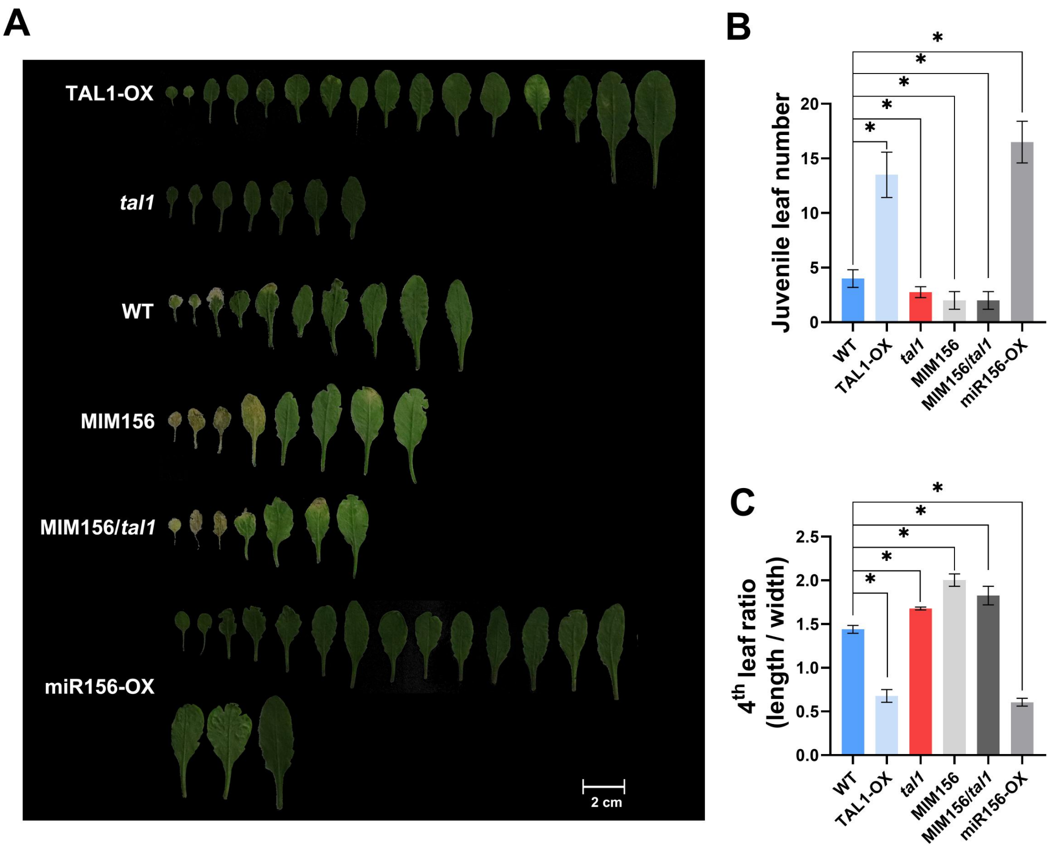


**Fig. S7 The aging phenotypes of TAL1-related lines in Arabidopsis at LD conditions.** The aging phenotypes of TAL1-OX, *tal1*, WT, MIM156 (mimic miR156), MIM156/ *tal1* (crossed line of MIM156 and *tal1*), and miR156-OX (miR156 overexpression line) were presented by all leaves in long day condition (A) and compered by their juvenile leaf number (B) and 4$^{th}$ leaf ratio (C). The '*' represents significant differences between samples at $P < 0.05$, based on a one-way ANOVA, with multiple comparisons made using Tukey's test.

### Figure S8

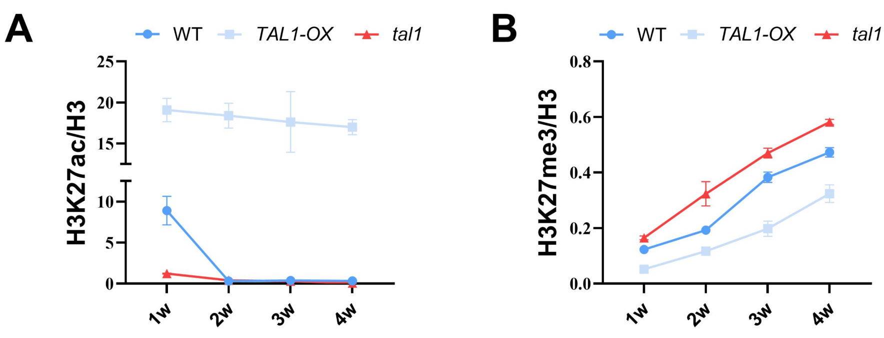


**Fig. S8 TAL1 act to promote acetylation of H3K27ac and suppress methylation of H3K27me3.** Dynamics of H3K27ac (A) and H3K27me3 (B) level of promoter region of MIR156A were measured by ChIP-qPCR.

### Figure S9

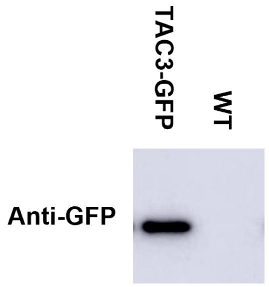


**Fig. S9 Identification of TAC3 overexpression line in Arabidopsis.** TAC3-GFP overexpression line were identified by Western Blot.

### Figure S10

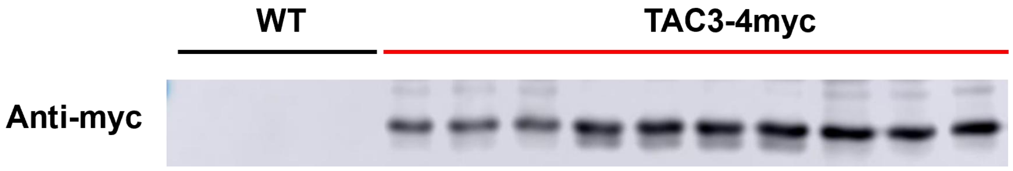


**Fig. S10 Identification of TAC3 overexpression line in Chinese fir.** TAC3-4MYC overexpression line were identified in Chinese fir by Western Blot.

### Figure S11

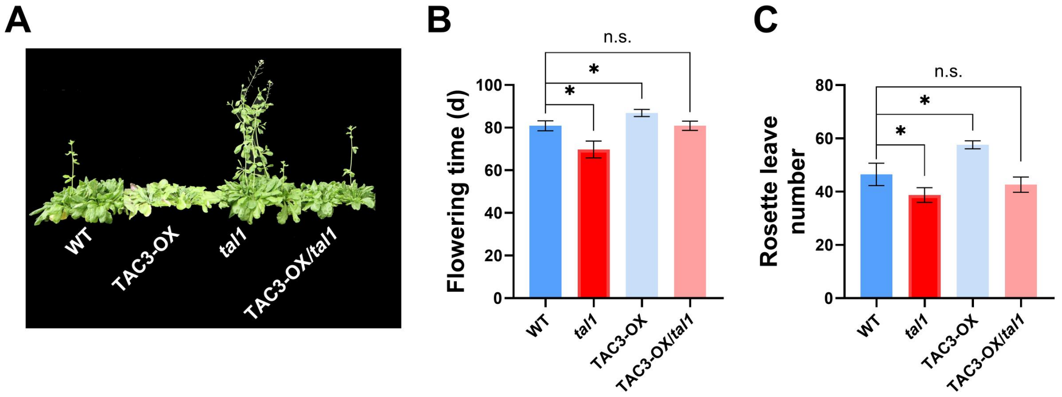


**Fig. S11 TAC3 suppress flowering in Arabidopsis at SD conditions** The function of TAC3 was explored in Arabidopsis, the flowering phenotypes (A) of WT (wild type), TAC3 overexpression line (TAC3-OX), TAL1 single mutation line (*tal1*) and TAC3-OX/ *tal1* (crossed line of TAC3-OX and *tal1*) were compered by their flowering time (B), rosette leave number (C). The '*' and 'n.s.' represents significant differences between samples at $P < 0.05$ and $P > 0.05$, respectively, based on a one-way ANOVA, with multiple comparisons made using Tukey's test.

### Figure S12

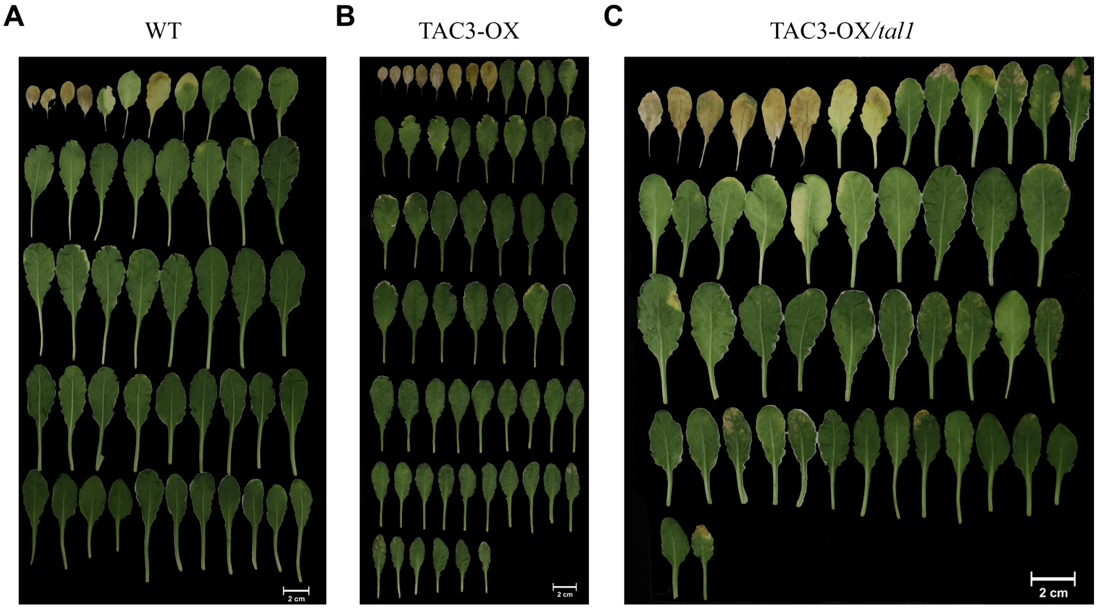


**Fig. S12 The aging phenotypes of TAC3-related lines in Arabidopsis at SD conditions**. The aging phenotypes of WT (A), TAC3-OX (B), and TAC3-OX/ *tal1* (crossed line of TAC3-OX and *tal1*) (C) were presented by all leaves in short day condition.

### Figure S13

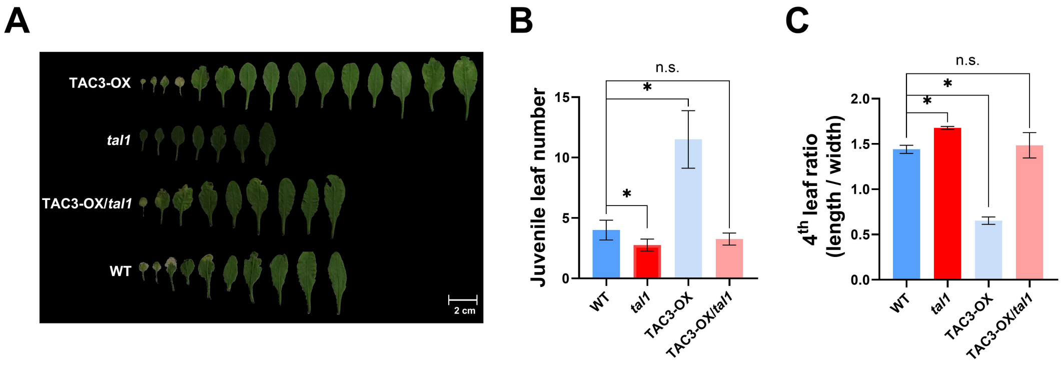


**Fig. S13 The aging phenotypes of TAC3-related lines in Arabidopsis at LD conditions.** The aging phenotypes of WT, TAC3-OX, and TAC3-OX/ *tal1* (crossed line of TAC3-OX and *tal1*) were presented by all leaves in long day condition (A) and compered by their juvenile leaf number (B) and 4$^{th}$ leaf ratio (C). The '*' and 'n.s.' represents significant differences between samples at $P < 0.05$ and $P>0.05$, respectively, based on a one-way ANOVA, with multiple comparisons made using Tukey's test.

### Figure S14

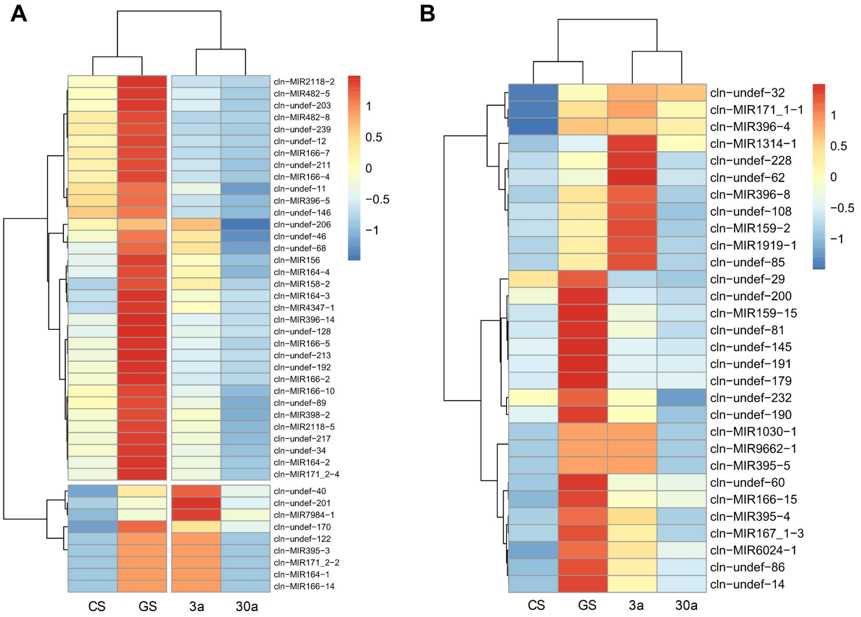


**Fig. S14. miRNA-seq analysis of different samples in Chinese fir.** Heatmaps were used to plot the expression levels of up- (A) and down-regulated (B) relevant genes by GS/CS and 3a/30a, respectively. Each row represents one gene, each column represents one sample, and the data results are homogenized by row. The color ranges from blue to red, indicating FPKM from small to large.

### Figure S15

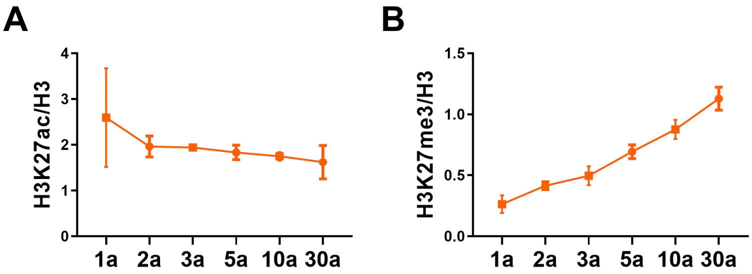


**Fig. S15 Dynamics of H3K27ac and H3K27me3 level in different age Chinese fir.** Dynamics of H3K27ac (A) and H3K27me3 (B) level of promoter region of MIR156A were measured in different age Chinese fir (1a, 2a, 3a, 5a, 10a and 30a) by ChIP-qPCR. Values were normalized by that of 1w.

#### Figure S16

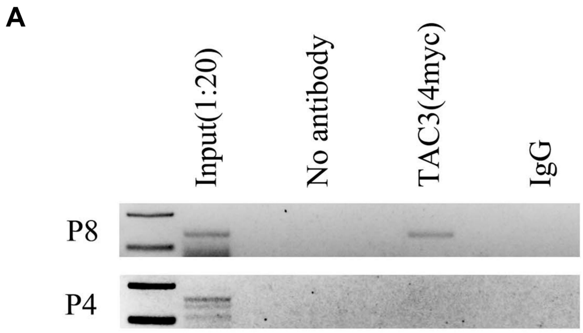


**Fig. S16 PCR analysis of P4 (negative control) and P8 in ChIP of Chinese fir.**

#### Figure S17

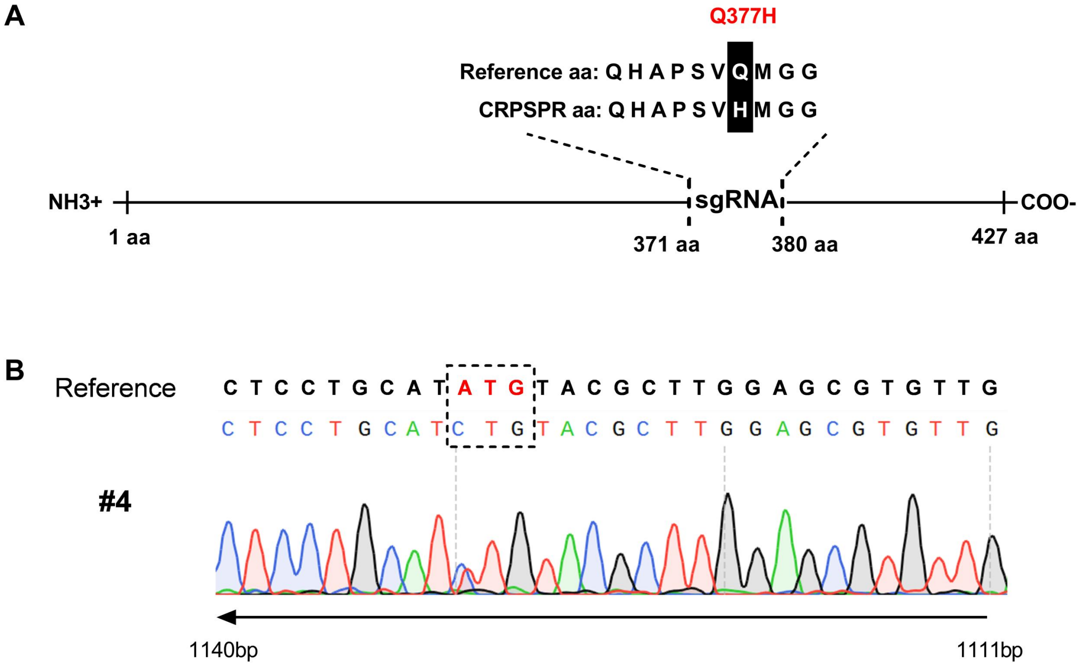


**Fig. S17 Cas9-AtHDA9-GFP edited the genome efficiently.** (A) AtHDA9 sgRNA sequences and Q377H mutant are shown in full CDS region. (B) Sanger sequencing results of representative T1 transgenic lines. BRI1 sgRNA sequences are underlined, and protospacer-adjacent motif (PAM) sequences are labeled in red.

Figure S18

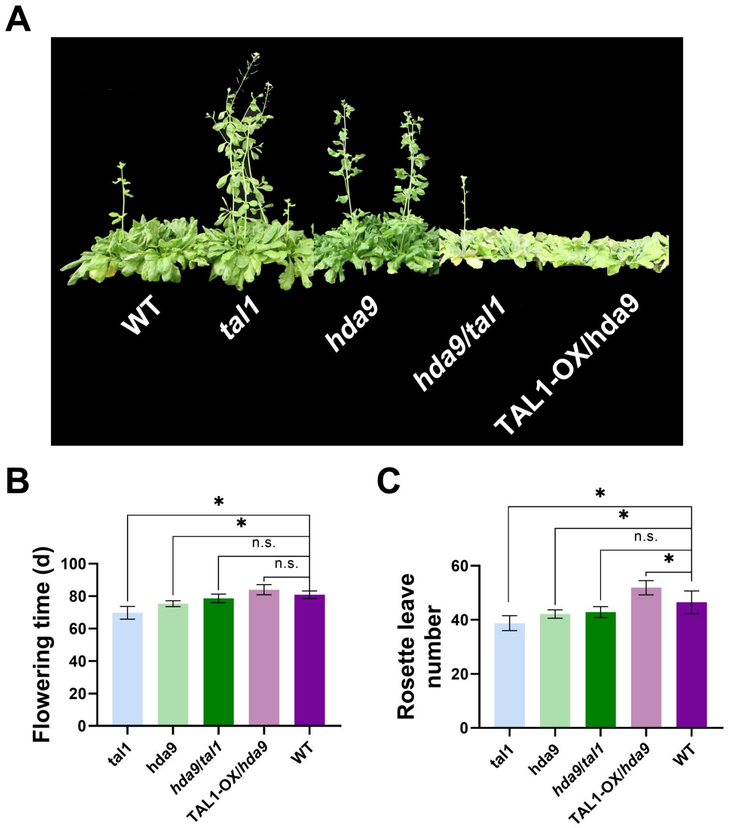


**Fig. S18 TAL1 suppress flowering through HDA9 in Arabidopsis at SD conditions.** The flowering (A) of WT (wild type), TAL1 single mutation line (*tal1*), HDA9 single(*hda9*), double (*hda9tal1*) mutation line and TAL1-OX/ *hda9* (crossed line of TAL1-OX and *hda9*) were compered by their flowering time (B), rosette leave number (C). The '*' and 'n.s.' represents significant differences between samples at $P < 0.05$ and $P>0.05$, respectively, based on a one-way ANOVA, with multiple comparisons made using Tukey's test.

### Figure S19

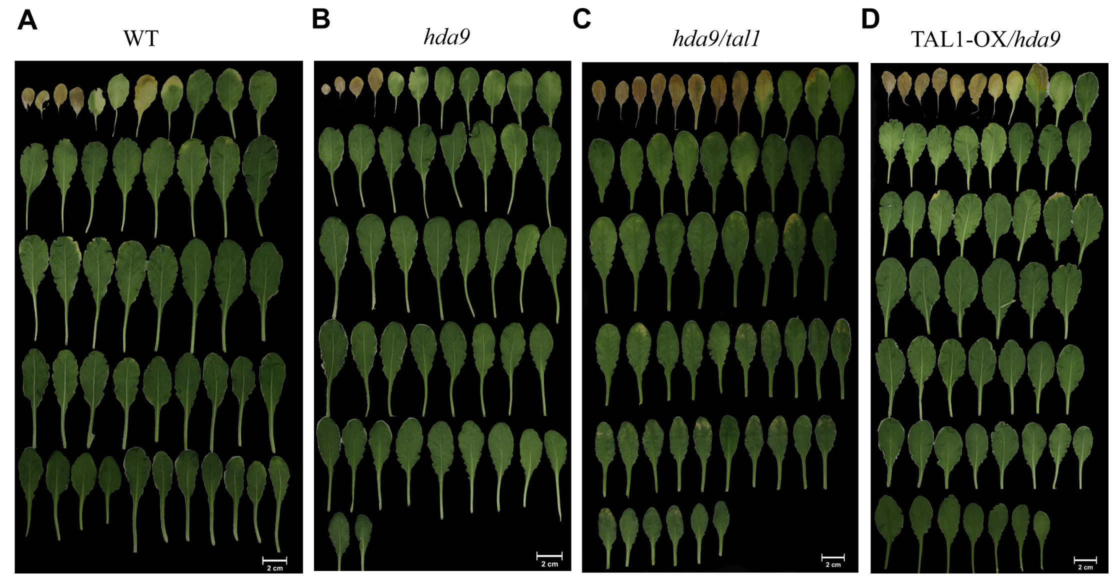


**Fig. S19 The aging phenotypes of TAL1 and HDA9-related in Arabidopsis at SD conditions**. The aging phenotypes of WT (A), *hda9* (B), *hda9/ tal1* (crossed line of *hda9* and *tal1*) (C) and TAL1-OX/*hda9* (crossed line of TAL1-OX and *hda9*) (D) were presented by all leaves in short day condition.

### Figure S20

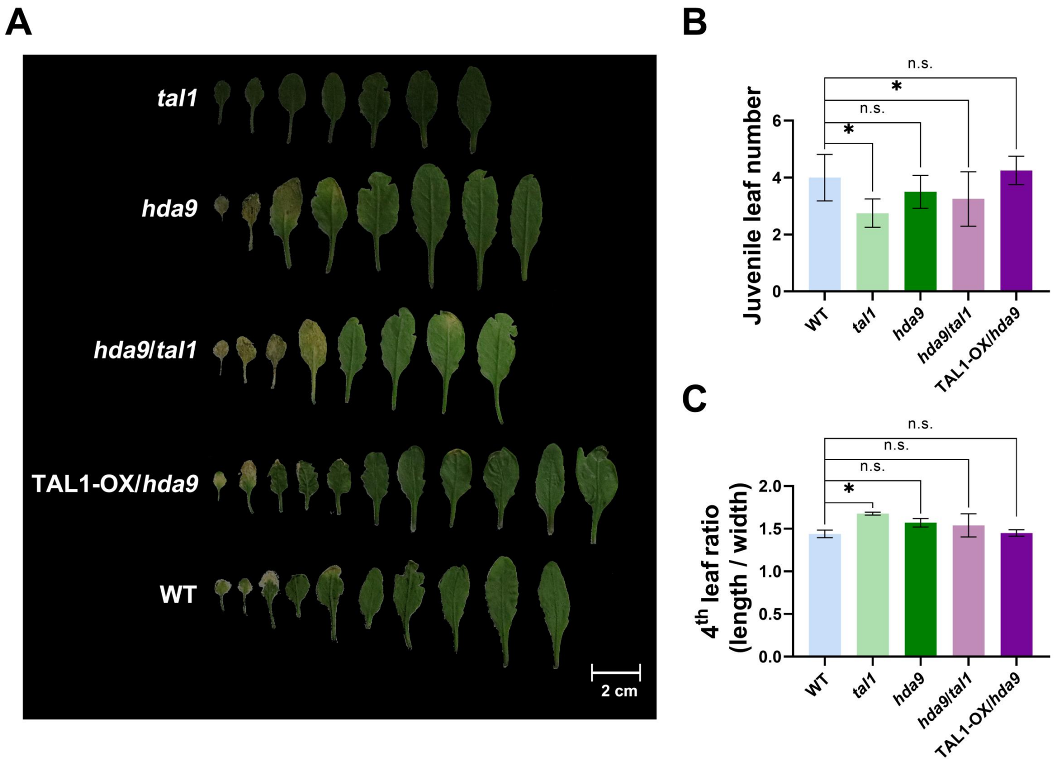


**Fig. S20 The aging phenotypes of TAL1 and HDA9-related lines in Arabidopsis at LD conditions.** The aging phenotypes (A) of WT, *tal1*, *hda9*, *hda9/ tal1* (crossed line of *hda9* and *tal1*) and TAL1-OX/*hda9* (crossed line of

TAL1-OX and *hda9*) were presented by all leaves in long day condition and compered by their juvenile leaf number (B) and 4$^{th}$ leaf ratio (C). The '*' and 'n.s.' represents significant differences between samples at P < 0.05 and P>0.05, respectively, based on a one-way ANOVA, with multiple comparisons made using Tukey's test.

### Figure S21

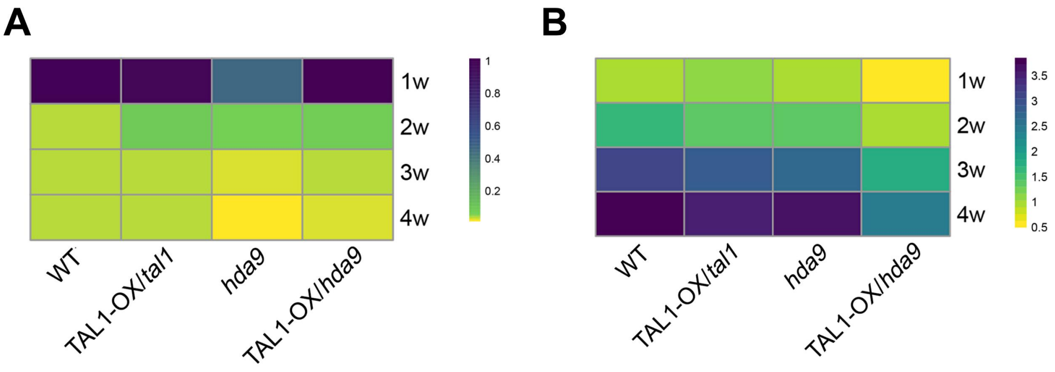


**Fig. S21 TAL1 and HDA9 act together to regulated acetylation of H3K27ac and methylation of H3K27me3.** Dynamics of H3K27ac (A) and H3K27me3 (B) level of promoter region of MIR156A were measured by ChIP-qPCR. Values were normalized by that of 1w.

### Figure S22

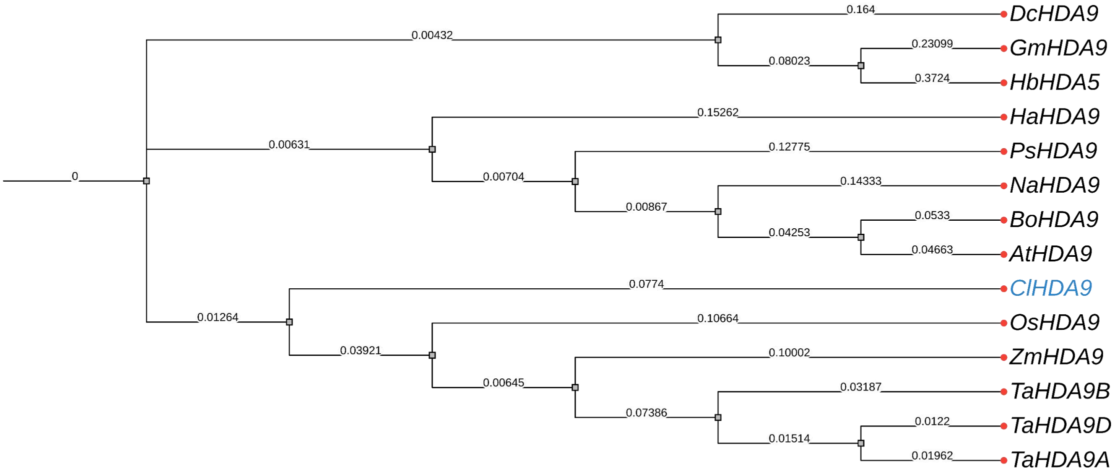


**Fig. S22 Phylogenetic analysis of ClHDA9 proteins in different plant species.** The phylogenetic tree was constructed based on the amino acid sequence of HDA9 and its orthologs from *Cunninghamia lanceolata* (Cl), *Pterocarya stenoptera* (Ps), *Arabidopsis thaliana* (At), *Oryza sativa* (Os), *Glycine max* (Gm), *Brassica oleracea* (Bo), *Nicotiana attenuate* (Na), *Hevea brasiliensis* (Hb), *Helianthus annuus* (Ha), *Dendrobium catenatum* (Dc), *Triticum aestivum* (Ta) and *Zea mays* (Zm). The accession numbers of the protein sequences are: Cula0010960.1 (ClHDA9), EU966459.1 (ZmHDA9), AT3G44680 (AtHDA9), MH444363.1 (BoHDA9), 100786705 (GmHDA9), XM_019382979.1 (NaHDA9), XM_021801905.2 (HbHDA9), XM_022138587.2 (HbHDA9), XM_028693057.1 (HaHDA9), XM_028693057.1 (DcHDA9), XM_044601483.1 (TaHDA9A), XM_044468425.1 (TaHDA9B), XM_044476676.1 (TaHDA9D), XM_015780381.1 (OsHDA9), XM_011007018.1 (PsHDA9). The phylogenetic tree was constructed from protein sequences obtained from the National Center for Biotechnology Information

(NCBI), and the analysis was performed as described previously and modified online in iTOL (https://itol.embl.de/).

### Figure S23

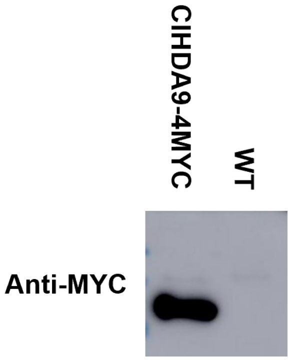


**Fig. S23 Identification of ClHDA9 overexpression line in Arabidopsis.** ClHDA9-4MYC overexpression line were identified by Western Blot.

### Figure S24

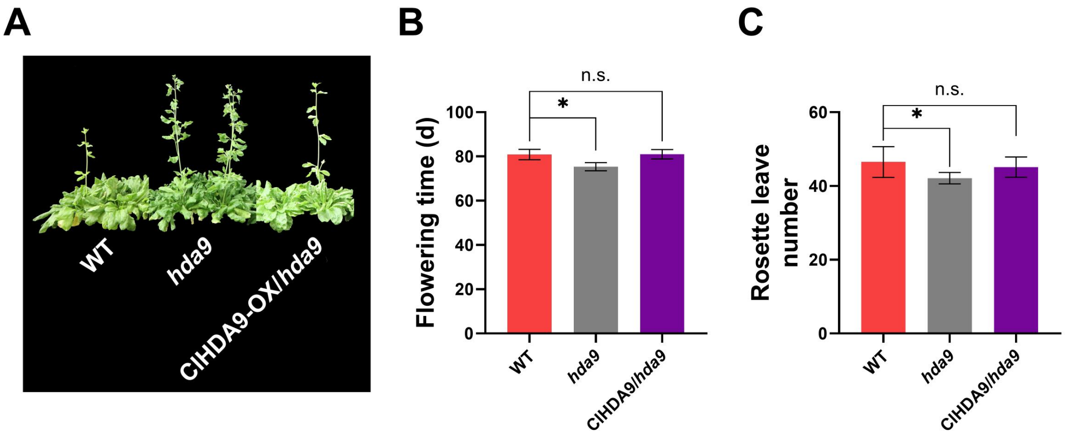


**Fig. S24 HDA9 suppress flowering in Arabidopsis at SD conditions.** The flowering (A) of WT (wild type), HDA9 single(*hda9*) and ClHDA9-OX/ *hda9* (crossed line of ClHDA9-OX and *hda9*) were compered by their flowering time (B), rosette leave number (C). The '*' and 'n.s.' represents significant differences between samples at $P < 0.05$ and $P>0.05$, respectively, based on a one-way ANOVA, with multiple comparisons made using Tukey's test.

### Figure S25

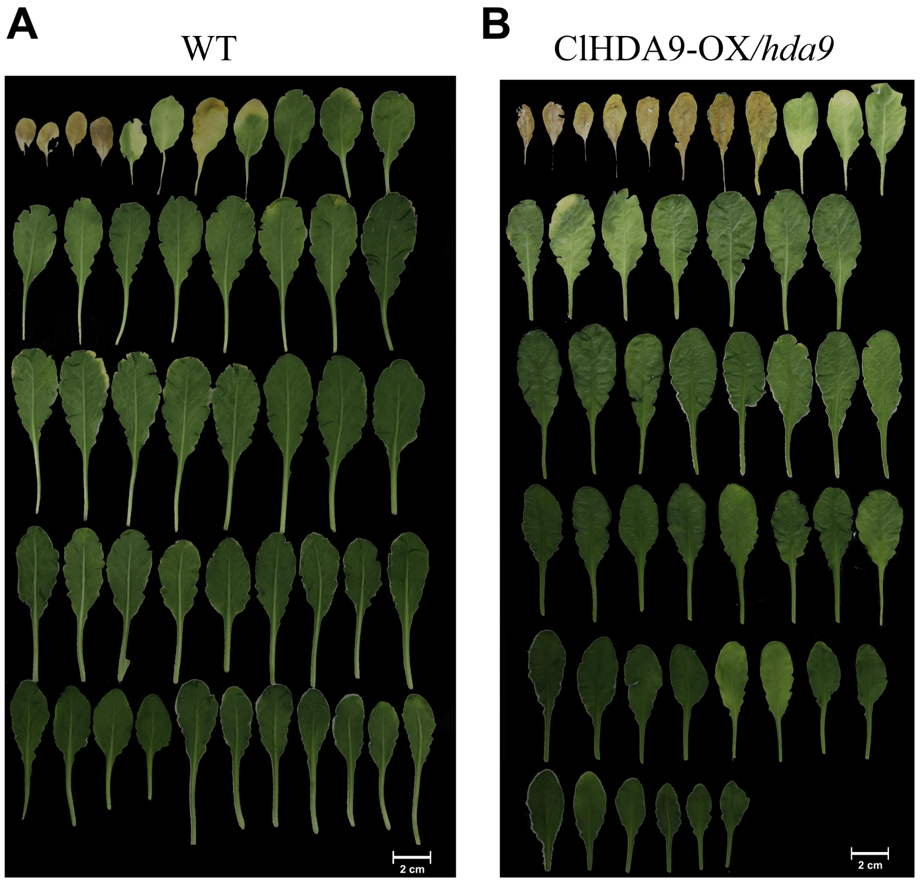


**Fig. S25 The aging phenotypes of ClHDA9-related in Arabidopsis at SD conditions**. The aging phenotypes of WT (A) and ClHDA9-OX/ *hda9* (crossed line of ClHDA9-OX and *hda9*) (B) were presented by all leaves in short day condition.

### Figure S26

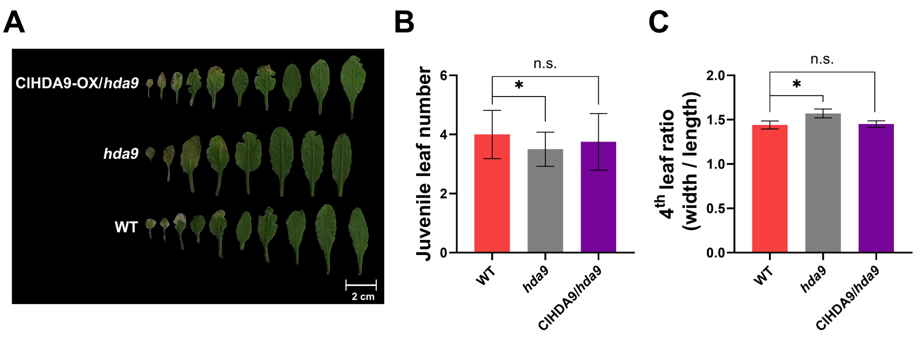


**Fig. S25 The aging phenotypes of ClHDA9-related lines in Arabidopsis at LD conditions.** The aging phenotypes of WT (wild type), HDA9 single(*hda9*) and ClHDA9-OX/ *hda9* (crossed line of ClHDA9-OX and *hda9*) were presented by all leaves in long day condition (A) and compered by their juvenile leaf number (B) and

4th leaf ratio (C). The '*' and 'n.s.' represents significant differences between samples at $P < 0.05$ and $P>0.05$, respectively, based on a one-way ANOVA, with multiple comparisons made using Tukey's test.

### Figure S26

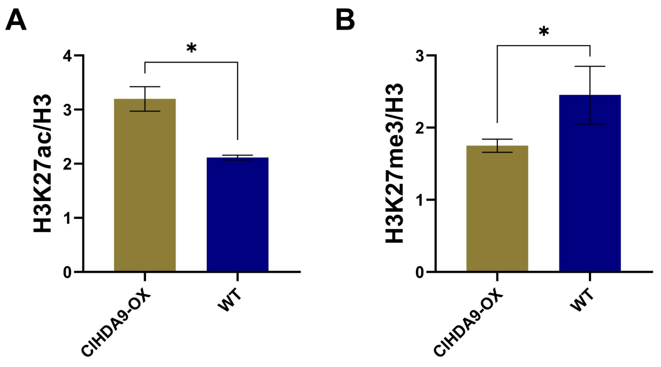


**Fig. S26 ClHDA9 act to suppress acetylation of H3K27ac and promote methylation of H3K27me3 in Chinese fir.** Dynamics of H3K27ac (A) and H3K27me3 (B) level of promoter region of MIR156A were measured in Chinese fir by ChIP-qPCR.
